# Cysteine-rich receptor-like kinases mediate Wall Teichoic Acid perception in *Arabidopsis*

**DOI:** 10.1101/2025.09.18.677107

**Authors:** L. Pierdzig, C. Seinsche, S. Kalla, S. Armiento, L.M. Schulz, J. Rismondo, A. Molinaro, C. De Castro, E.K. Petutschnig, V. Lipka

## Abstract

Plants and animals detect microbe-associated molecular patterns (MAMPs) to initiate defense responses. Both lineages employ pattern recognition receptors (PRRs), yet plant mechanisms for sensing Gram-positive bacterial MAMPs remain poorly understood. Wall teichoic acids (WTAs), anionic glycopolymers unique to Gram-positive bacteria, are shown here to activate salicylic acid signaling, defense priming, and a non-canonical programmed cell death in *Arabidopsis*. We demonstrate that glycosylated WTAs from diverse Gram-positive species elicit these responses and provide genetic evidence that their recognition depends on two members of the plant-specific CYSTEINE-RICH RECEPTOR-LIKE KINASE (CRK) gene family, which is absent in animals and whose function remains elusive. These findings reveal WTAs as a novel class of MAMPs in plants and highlight CRKs as key components in their perception.

## Introduction

Animals and plants are continuously exposed to a wide array of microbial pathogens, commensals, and symbionts. A fundamental feature of innate immunity in both kingdoms is the recognition of Microbe-Associated Molecular Patterns (MAMPs) by pattern recognition receptors (PRRs) (*1, 2*). In plants, PRRs predominantly comprise cell surface receptors (CSRs), including receptor-like kinases (RLKs) and receptor-like proteins (RLPs), which operate in concert with membrane-associated co-receptors and receptor-like cytoplasmic kinases (RLCKs) to detect non-self ligands and to initiate signaling cascades that activate basal defense responses (*2, 3*). Plants possess multiple families of RLPs and RLKs, which vary considerably in size and composition across plant species (*4*). They are hypothesized to have expanded over evolutionary time under the selective pressures imposed by pathogen challenges (*4*). As a result, plant lineages exhibit extensive diversification of CSRs, which are classified based on their receptor domain architecture. LEUCINE-RICH REPEAT (LRR)-RLKs typically recognize proteinaceous ligands, such as bacterial flagellin or elongation factor Tu (EF-Tu), whereas LECTIN-RLKs (LecRLKs), LYSIN-MOTIF (LysM)-RLKs, MALECTIN-LIKE RLKs, and WALL-ASSOCIATED KINASE (WAK)-LIKE RLKs primarily detect carbohydrate-derived MAMPs (*4*). MAMP recognition by CSRs triggers a canonical set of downstream immune responses, including cytosolic calcium (Ca²⁺) influx, apoplastic alkalinization, reactive oxygen species (ROS) production, phosphorylation of downstream RLCKs, activation of the MITOGEN-ACTIVATED PROTEIN KINASE (MAPK) cascade, and transcriptional induction of defense-related genes. Collectively, these events constitute pattern-triggered immunity (PTI), forming a robust basal defense mechanism against microbial invasion (*5*).

In animals, innate immune surveillance is primarily mediated by intracellular PRRs, and the diversity of CSR classes is comparatively limited (*1*). These CSRs are classified based on their receptor domain architecture into two major groups: TOLL-LIKE RECEPTORS (TLRs) and C-TYPE LECTIN RECEPTORS (CLRs) (*1*). In vertebrates, particularly mammals, TOLL-LIKE RECEPTOR 2 (TLR2) recognizes lipoteichoic acid (LTA), a key structural component of Gram-positive bacterial cell walls (*6, 7*). This recognition event initiates canonical innate immune signaling pathways, facilitating host defense against Gram-positive infections and contributing to the regulation of inflammatory responses (*7*). Notably, while numerous MAMPs originating from Gram-negative bacteria have been identified in plants, elicitors of PTI that are exclusively associated with Gram-positive bacteria have, to date, not been characterized. This is unexpected, as Gram-positive bacteria are not only ancestral to Gram-negative bacteria (*8*) and extensively characterized as animal probiotics and pathogens (*9, 10*), but also represent abundant and ecologically important members of the plant rhizosphere and phyllosphere microbiomes (*11*). They can function as potent plant growth promoters, stimulators of plant symbioses, or phytopathogens (*11*). Here, we asked whether Gram-positive bacteria produce lineage-specific MAMPs with immune-activating properties and, if so, how these are recognized by the plant immune system.

## Results

### Gram-positive bacteria trigger cell death in *Arabidopsis*

Infiltration assays using the Gram-negative plant pathogen *Pseudomonas syringae* pv. *tomato* DC3000 (*Pst* DC3000) and the Gram-positive model strain *Bacillus subtilis* 168 revealed that both organisms can elicit disease-like symptoms in *Arabidopsis* leaves at 5 dpi (Fig. 1A). *B. subtilis* 168 triggered lesion formation only at high inoculum (10⁷ cfu/ml), whereas *Pst* DC3000 caused classical disease symptoms at both high and low titers. As expected, *Pst* DC3000 symptom development correlated with *in planta* growth, while *B. subtilis* 168 did not proliferate (Fig. 1B). Remarkably, infiltration of autoclaved *B. subtilis* 168, but not heat-killed *Pst* DC3000, induced similar lesions (Fig. 1A), suggesting that *B. subtilis* triggers leaf-specific cell death via a heat-stable factor or MAMP. To determine whether this activity is specific to *B. subtilis* or a general property of Gram-positive bacteria, we infiltrated *Arabidopsis* leaves with heat-killed cultures from multiple Gram-positive and Gram-negative species (Fig. 1C). Several Gram-positive bacteria, including the human pathogens *Staphylococcus aureus*, *Listeria monocytogenes* and the plant-associated *Clavibacter michiganensis* and *Lactoplantibacillus plantarum*, reproduced the lesion phenotype, whereas none of the Gram-negative strains tested, including the model species *Escherichia coli* and the plant pathogens *P. syringae*, *Agrobacterium tumefaciens*, and *Burkholderia plantarii*, induced comparable responses (Fig. 1C). These results point to a conserved cell death–inducing agent (CDA) present in Gram-positive, but absent in Gram-negative, bacteria.

**Fig. 1.**
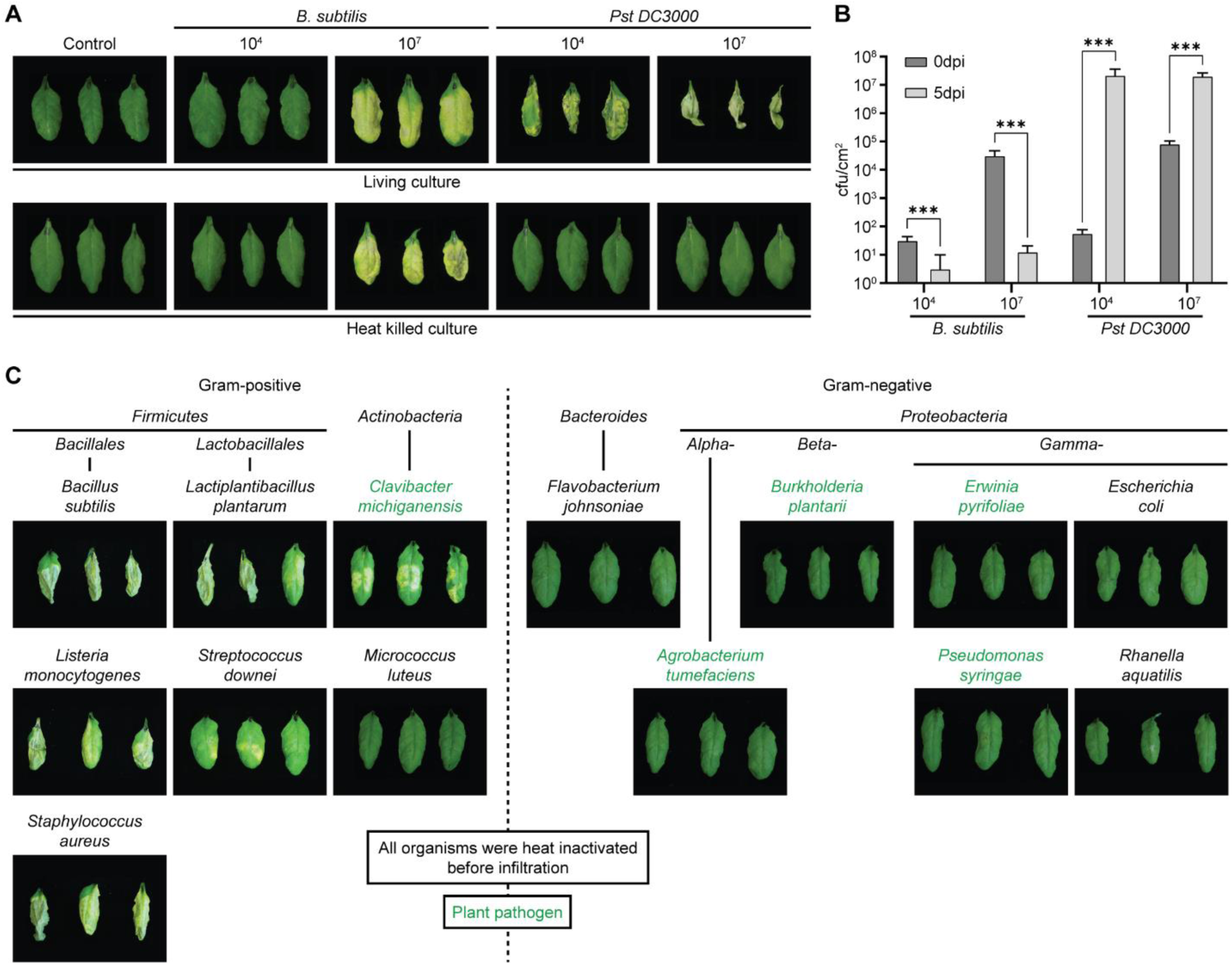
Heat-inactivated Gram-positive bacteria induce a macroscopically detectable lesion phenotype in *Arabidopsis* leaves. (**A**) Infiltration of 5-week-old *Arabidopsis* leaves with living and heat-inactivated cultures of *Pseudomonas syringae* pv *tomato* DC3000 (*Pst* DC3000) and *B*. *subtilis* 168 pBQ200 at 10^4^ or 10^7^ cfu/ml in infiltration medium (10 mM MgCl_2_, 0.0002% Silvert-L77). Infiltration medium alone was infiltrated as a control. Pictures were taken at 5 dpi. (**B**) Bacterial proliferation inside *Arabidopsis* leaves was determined by quantifying cfu/cm^2^ of infiltrated leaves at 0 and 5 dpi. Three leaf discs from three individual plants were pooled as one biological replicate. Bars represent mean cfu/cm^2^ leaf area ± SD (n=6). Asterisks indicate significant differences between 0 and 5 dpi time points (Student’s T-test, ** p<0.01, *** p<0.001). (**C**) Infiltration of 5-week-old *Arabidopsis* plants with different heat-inactivated Gram-positive and Gram-negative bacteria cultures with an OD_600_ of 0.2 in 10mM MgCl_2_. (A and C) Images represent three leaves from three independent plants.

### Glycosylated wall teichoic acid is the cell death-inducing agent in Gram-positive bacteria

Previous reports showed that lipopeptides from plant beneficial *Bacillus* strains such as surfactin and fengomycin can trigger induced systemic resistance in plants (*12, 13*). However, *B. subtilis* 168 lacks functional lipopeptide biosynthesis due to a *SURFACTIN PRODUCING* (*sfp*) frameshift mutation (*14, 15*), and exogenous surfactin failed to induce PCD (fig. S1), excluding lipopeptides as the CDA. Biochemical fractionation of heat-inactivated *B. subtilis* 168 cultures (fig. S2A) revealed that the CDA is heat-stable (121 °C), and water-soluble, as it is retained after ultracentrifugation (fig. S2, A and B). Proteinase K treatment had no effect, indicating a non-proteinaceous nature (fig. S2, C and D). Concanavalin A binding reduced the cell death phenotype (fig. S2E), suggesting glycan components, and ion exchange chromatography identified the CDA as anionic (fig. S2, F and G).

Given these features, we investigated wall teichoic acids (WTAs) as possible CDA. WTAs are abundant and surface-exposed anionic glycopolymers of Gram-positive cell walls that are covalently bound to the peptidoglycan layer (fig. S3) (*16, 17*). Recent findings have shown that natural diversity of *B. subtilis* is accompanied by variations in WTA biosynthesis (*18*). Analysis of diverse *B. subtilis* isolates with defined WTA compositions (*18*) revealed that strains predicted to produce glycerol-phosphate (GroP) WTAs (168, BSn5, BEST195, BEST7003) strongly induced cell death, whereas a ribitol-phosphate (RboP) WTA strain (W23) showed weaker activity, and a WTA-deficient strain (TLO3) failed to elicit responses (fig. S4, B and C).

Targeted mutants of WTA biosynthesis or glycosylation in *B. subtilis* 168, and the two human pathogen strains *S. aureus* NRS384, and *L. monocytogenes* 10403S lost cell death–inducing capacity (Fig. 2, A and B). Notably, wildtype *S. aureus* NRS384 and *L. monocytogenes* 10403S, which both produce RboP-based WTAs (*19, 20*), showed weaker cell death activity compared to the GroP-based WTA producing *B. subtilis* 168 (*16*) (Fig. 2B). Thus, GroP-based WTAs were generally more active than RboP-based WTAs (Fig. 2B and fig. S4C), suggesting either multiple recognition pathways or differential affinity for a single receptor.

**Fig. 2.**
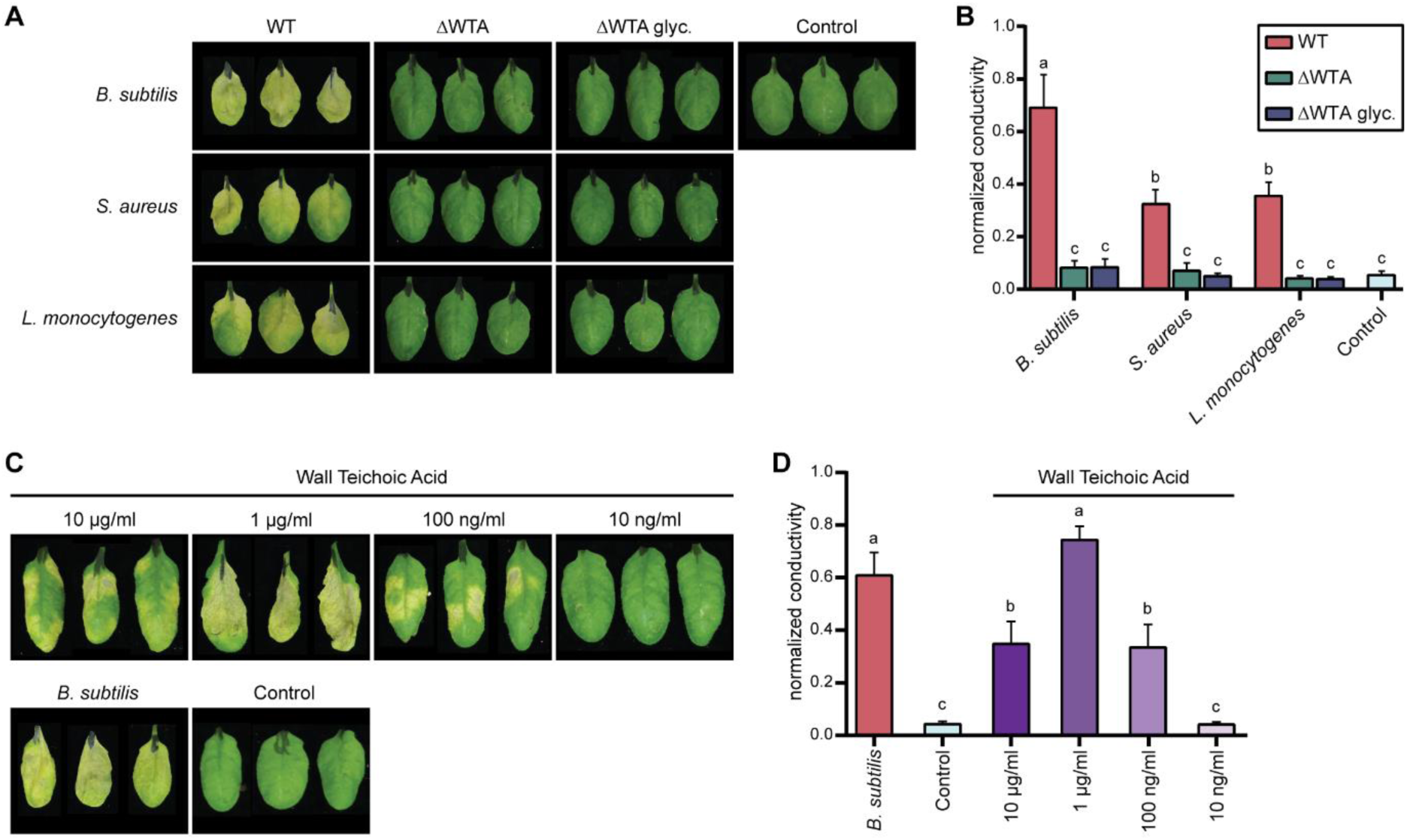
Glycosylated Wall Teichoic Acid (WTA) is the cell death inducing agent in *Bacillus subtilis* extracts. (**A** and **B**) Infiltration of 5-week-old *Arabidopsis* Col-0 plants with heat-inactivated *B*. *subtilis* 168, *Staphylococcus aureus* NRS384, and *Listeria monocytogenes* 10403S extracts. For each species, a wild-type strain, a WTA biosynthesis mutant (ΔWTA), and a WTA glycosylation mutant (ΔWTA glyc.) were infiltrated at an OD_600_ of 0.02 in 10 mM MgCl_2_. 10 mM MgCl_2_ was infiltrated as a control. (**C** and **D**) Infiltration of 5-week-old *Arabidopsis* Col-0 with pure WTAs purified from *B*. *subtilis* 168 at decreasing concentrations from 10 µg/ml to 10 ng/ml. Heat-inactivated *B*. *subtilis* 168 extract and 10 mM MgCl_2_ were infiltrated as positive and negative controls, respectively. (A and D) Pictures show three infiltrated leaves from three individual plants at 6 dpi. (B and C) Ion leakage assay to quantify cell death at 6 dpi. Bars represent mean normalized conductivity ± SEM (n=12). Different letters indicate significant differences of means (One-way ANOVA with Tukey’s post hoc test, p<0.05).

Purification of *B. subtilis* 168 WTAs via size-exclusion and anion exchange chromatography followed by 2D NMR identified glycosylated GroP-WTA as the active molecule (fig. S5, A to D). Infiltration of purified WTA induced a dose-dependent cell death response in *Arabidopsis* at concentrations from 100 ng/ml to 10 µg/ml (Fig. 2, C and D). WTAs isolated via an established extraction protocol (*21*) produced similar activity (fig. S6). Lipoteichoic acid (LTA), a structural analog of WTA with an enantiomeric backbone that is attached to the bacterial plasma membrane via a glycolipid anchor (fig. S3) (*22, 23*), is a known Gram-positive MAMP in vertebrates (*7*) and elicits nitric oxide (NO) production in *Arabidopsis* (*24*). However, a *B. subtilis* 168 Δ*ltaS*Δ*yfnI*Δ*yqgS*Δ*yvgJ* mutant deficient in LTA biosynthesis (*25*) retained full cell death-inducing activity (fig. S7, A and B). This indicates that, in contrast to vertebrates, where both LTA and WTA act as MAMPs (*7, 26*), plants predominantly perceive glycosylated WTA as a Gram-positive–specific elicitor.

### WTA-induced programmed cell death requires CYSTEINE-RICH RECEPTOR-LIKE KINASE 7 (CRK7)

We next asked whether the cell death triggered by WTA reflects programmed cell death (PCD) initiated by WTA perception. Mutations in canonical *Arabidopsis* PRRs, including FLAGELLIN SENSITIVE 2 (FLS2), CHITIN ELICITOR RECEPTOR KINASE 1 (CERK1), and LIPOOLIGOSACCHARIDE-SPECIFIC REDUCED ELICITATION (LORE), did not affect WTA-induced PCD (Fig. 3A). A forward genetic screen looking for plants with altered cell death responses yielded several *Arabidopsis* mutant lines that were insensitive to WTA, including the *crk7-4* mutant (fig. S8A). Mapping-by-sequencing localized the causal mutation to chromosome 4 (fig. S8B), revealing a non-synonymous single-nucleotide polymorphism (nsSNP) affecting the second conserved cysteine in the first DUF26 domain of CYSTEINE-RICH RECEPTOR-LIKE KINASE 7 (CRK7) (fig. S8, C and D). Additional T-DNA insertion lines (*crk7*-5, *crk7*-6, *crk7*-7, *crk7*-8) and mutants of closely related and adjacent CRK genes (*CRK6, CRK8*) confirmed that only loss of CRK7 suppressed WTA-induced cell death (fig. S8A). Transgenic complementation with wild-type or mCitrine-tagged CRK7 under its native promoter fully restored the phenotype (Fig. 3B and fig. S9).

**Fig. 3.**
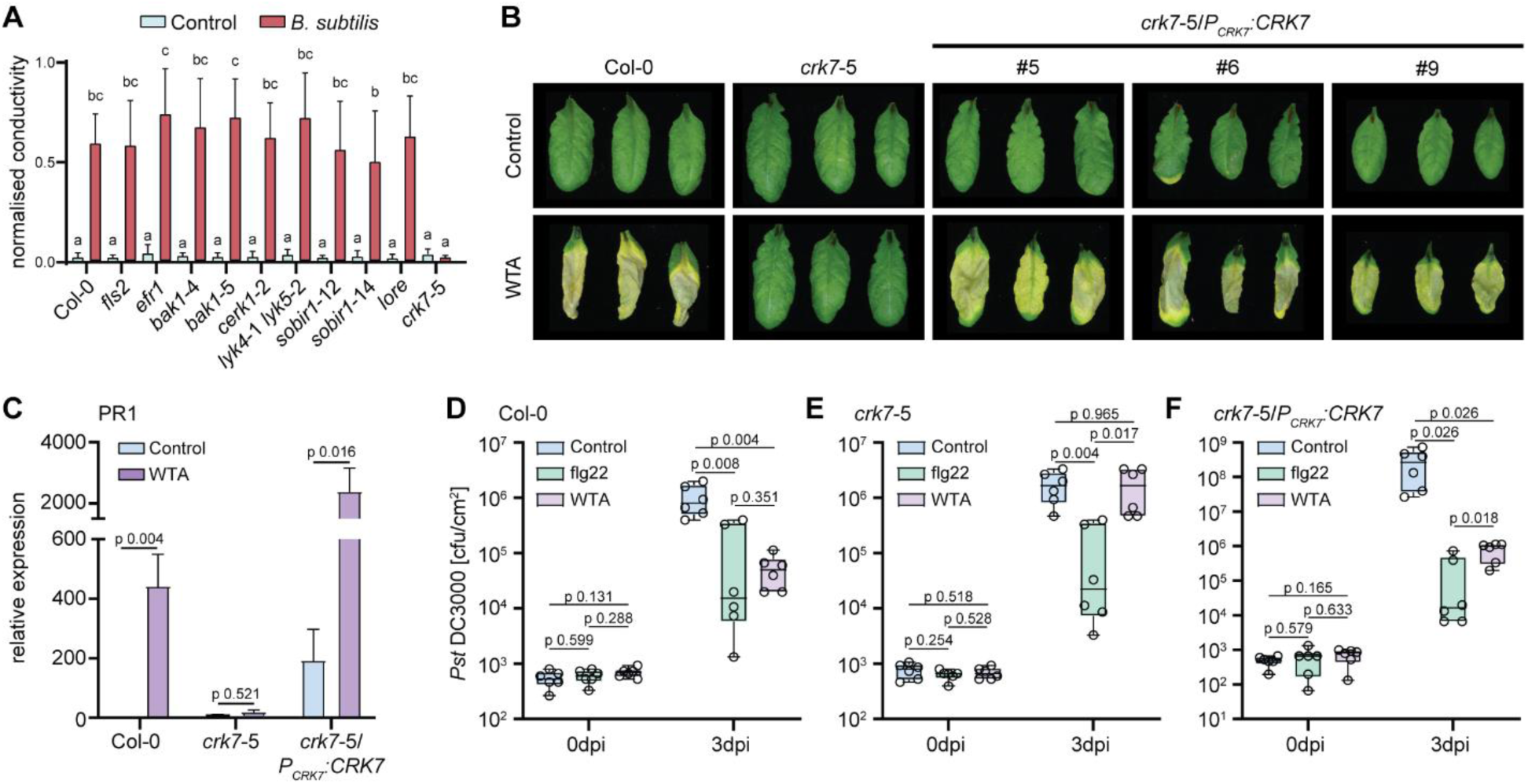
Cysteine-rich Receptor-like Kinase 7 (CRK7) is required for WTA-induced responses. (**A**) Ion leakage assay of different *Arabidopsis* PTI receptor mutants at 6 dpi. Leaves were infiltrated with heat-inactivated *B*. *subtilis* at an OD_600_ of 0.02 in 10 mM MgCl_2_. Bars represent mean normalized conductivity ± SEM (n=12). Different letters indicate significant differences of means (One-way ANOVA with Tukey’s post hoc test, p<0.05). (**B**) Images of 5-week-old *Arabidopsis* Col-0, *crk7*-5, and 3 individual *crk7*-5 lines complemented with genomic CRK7 expressed under its native promoter (*crk7*-5/*P_CRK7_:CRK7*) at 6 dpi. (**C**) qRT-PCR to monitor PR1 expression in response to WTA treatment in Col-0, *crk7*-5, and *crk7*-5/*P_CRK7_:CRK7*. Bars represent mean relative PR1 expression ± SEM (n>5 biological replicates, Student’s T-test between WTA and control treatment). (B and C) Leaves were infiltrated with 1 µg/ml WTA or 10 mM MgCl_2_ as a control. (**D** to **F**) Leaves of Col-0, *crk7*-5, and *crk7*-5/*P_CRK7_:CRK7* were pretreated for 24 h with 10 ng/ml WTA, 10 mM MgCl_2,_ or 100 nM flg22. Pretreated leaves were infiltrated with *Pst* DC3000. Colony-forming units per cm^2^ leaf area (CFU/cm^2^) were counted at 0 and 3 dpi. Bounds of boxplots represent the 25^th^ and 75^th^ percentile; the line shows the median, and the whiskers the 1.5xIQR (n=6; Welch’s T-test).

### CRK7 mediates priming and WTA-triggered defense gene expression

WTA did not induce canonical pattern-triggered immunity (PTI) outputs, such as apoplastic ROS or Ca²⁺ influx (fig. S10), but did trigger CRK7-dependent expression of the salicylic acid (SA) marker gene *PR1* at 2 dpi (Fig. 3C). Pre-treatment with sublethal concentrations of WTA (10 ng/µL) enhanced resistance against *Pst* DC3000, similar to 100 nM flg22 (Fig. 3D). This priming effect was abolished in the *crk7*-5 mutant and restored upon CRK7 complementation (Fig. 3, E and F). Gram-positive bacteria, including *Bacillus* spp., are common members of the phyllosphere and rhizosphere microbiota (*11, 27, 28*), suggesting that WTA perception may have an important immune-modulatory function and may contribute to microbiome-driven immune priming in *Arabidopsis*.

### CRK7-dependent WTA-induced cell death is confined to the genus *Arabidopsis*

Testing a range of monocot and dicot species revealed that WTA-induced PCD was confined to *Arabidopsis* species (*A. thaliana, A. suecica, A. shokei*) (fig. S11A), correlating with the presence of a CRK7 homolog (fig. S11B). To assess the natural variation in WTA responsiveness within *A. thaliana*, we screened 113 accessions for PCD following infiltration with pre-purified *B. subtilis* extracts. Of these, 92 accessions displayed robust cell death, whereas 21 were insensitive (fig. S12). Comparative sequence analysis of the published genomes of 108 (*29*) of the 113 accessions showed that all but one insensitive accession, Cvi-0, retained a CRK7 homolog. Cvi-0 carried a complete deletion of the CRK7 locus, while 13 of the remaining 15 insensitive lines encoded divergent CRK7 variants clustering together phylogenetically. Two insensitive accessions, Ws-0 and Gd-1, harbored CRK7 alleles closely related to those in responsive accessions, indicating that loss of sensitivity in these lines may involve defects in additional components of the WTA signaling pathway.

### CRK7 forms a preassembled complex with CRK43 to mediate WTA perception

Sequence comparison across insensitive accessions revealed multiple nsSNPs within CRK7, predominantly in the extracellular domain (fig. S13A). AlphaFold Multimer modeling (*30–31*) predicted CRK7 ectodomain homodimerization (fig. S13, B and C), which was confirmed by co-immunoprecipitation (Co-IP) and Fluorescence Lifetime Imaging–Förster Resonance Energy Transfer (FLIM-FRET) assays in *Nicotiana benthamiana* (Fig. 4, A to D). CRK7 homodimerized constitutively but underwent conformational changes upon WTA exposure, as reflected by reduced donor fluorescence lifetime (Fig. 4, B and C).

**Fig. 4.**
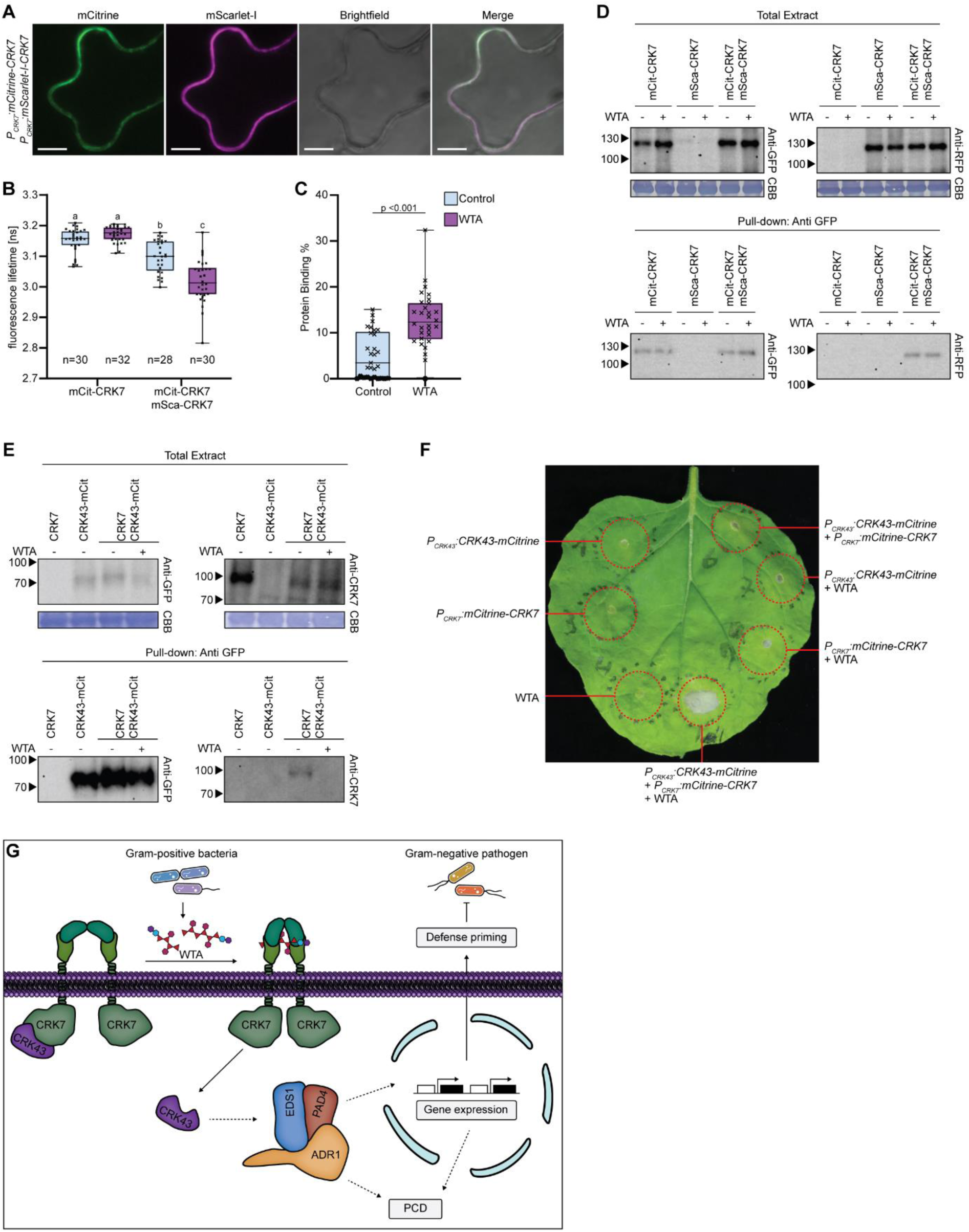
WTA controls CRK7 and CRK43 heteromer formation. (**A**) Confocal laser scanning microscopy of mCitrine- and mScarlet-I-tagged CRK7 transiently co-expressed from the native promoter in *N. benthamiana*. mCitrine and mScarlet-I were excited at 514 nm or 561 nm and detected at 525 nm to 555 nm or 580 nm to 620 nm, respectively. Scale Bar = 10 µm. (**B**) Fluorescence-lifetime imaging microscopy (FLIM) of mCitrine-tagged CRK7 alone and co-expressed with mScarlet-I-tagged CRK7, infiltrated with 1 µg/ml WTA (purple) or 10 mM MgCl_2_ (blue) as control 1 h before microscopy. The number of individual images (n) used for the analysis is indicated in the figure. Different letters indicate significant differences of means (One-way ANOVA with Tukey’s post hoc test, p<0.05). (**C**) Protein binding in [%] was calculated from FLIM-FRET analysis of mCitrine labeled CRK7 co-expressed with mScarlet-I labeled CRK7, treated with 1 µg/ml WTA or 10 mM MgCl_2_. (Student’s t-test, *** P<0.0001). (**D**) Anti-GFP pull-down experiment confirming the homomerization of CRK7 tagged with mCitrine and mScarlet-I. (**E**) Anti-GFP pulldown experiment demonstrating interaction between CRK7 and CRK43, which dissociates upon WTA treatment. (**F**) mCitrine-tagged CRK7 and CRK43 were transiently expressed in *N. benthamiana* alone, in combination, and together with 1 µg/ml WTA. Image was taken at 6 dpi (**G**). Current working model. CRK7 and CRK43 form a heteromeric complex at the plasma membrane. Upon WTA recognition, CRK43 dissociates from the complex to induce downstream signaling via the EDS1-PAD4-ADR1 signaling hub.

A second suppressor identified in our forward genetic screen, *crk43-1* (fig. S14A), carried a splice site mutation in another CYSTEINE-RICH RECEPTOR-LIKE KINASE, *CRK43*, leading to a premature stop codon (fig. S14, B to D). CRK43 represents one of three N-terminally truncated *CRKs* in the *Arabidopsis* genome, whose gene products apparently lack a ligand-binding and transmembrane domain, thus classifying them as soluble RLCKs (*33*). Both *crk43-1* and a T-DNA allele (*crk43*-2) failed to undergo WTA-induced PCD, and complementation restored the phenotype (fig. S14A). Co-IP experiments demonstrated constitutive CRK7–CRK43 interaction that dissociated upon WTA perception (Fig. 4E). Transient co-expression of CRK7 and CRK43 in *N. benthamiana*, which is normally unresponsive to WTA (fig. S11), reconstituted WTA-dependent PCD (Fig. 4F). Together, these data support a model in which CRK7 homodimers form a preassembled plasma membrane complex with cytosolic CRK43. Upon WTA recognition, CRK43 is released to initiate downstream signaling.

### WTA-induced cell death requires the EDS1–PAD4–ADR1 signaling node

Our forward genetic screen identified a third suppressor mutant carrying an nsSNP in *PHYTOALEXIN DEFICIENT 4 (PAD4)*, substituting Ser_499_ with Phe (S_499_F), hereafter referred to as *pad4-14* (fig. S15, A to D). Knockout (*pad4-1*) and *pad4-14* mutants both failed to undergo WTA-induced cell death (fig. S15, A and E), indicating that PAD4 is essential for this process and that the S_499_F mutation likely abolishes its function. PAD4, together with ENHANCED DISEASE SUSCEPTIBILITY 1 (EDS1) and SENESCENCE-ASSOCIATED GENE 101 (SAG101), constitutes the lipase-like EDS1 protein family, which functions as a central regulator of intracellular plant immunity and programmed cell death (*34*). Similar to animals, plants utilize cytosolic NUCLEOTIDE-BINDING LEUCINE-RICH REPEAT (NLR) receptors as part of their innate immune system (*1, 35*). However, unlike animals, plants have evolved distinct TOLL-INTERLEUKIN 1 RECEPTOR (TIR)-and COILED-COIL (CC)-type NLRs to detect pathogen-derived effectors within the cytoplasm (*35, 36*). Effector-triggered immunity (ETI) is mediated by mutually exclusive heterodimeric complexes of EDS1-SAG101 or EDS1-PAD4 (*34*). These heterodimers act as receptors for small signaling molecules, including phosphoribosyl AMP/ADP (pRIB-AMP/ADP) and ADP-ribosylated ATP (ADPr-ATP), generated by effector-activated TIR-NLR resistosomes (*37, 38*). Binding of ADPr-ATP to the EDS1-SAG101 complex or pRIB-AMP/ADP to the EDS1-PAD4 complex facilitates their association with specific CC-type helper NLRs (hNLRs), namely N REQUIRED GENE 1 (NRG1)-type and ACTIVATED DISEASE RESISTANCE 1 (ADR1)-type, respectively (*37, 38*). Notably, the EDS1–PAD4–ADR1 signaling module has recently been identified as a critical convergence node for defense signaling pathways initiated by both cell surface and intracellular LRR receptors (*34, 39*). Given that PAD4 is essential for WTA-induced cell death, we investigated whether this response is mediated via the EDS1–PAD4–ADR1 signaling module. To address this, WTA infiltration assays were conducted in *eds1*-2, *pad4-1*, *sag101*-3, *adr1* triple, and *nrg1* triple mutants. The assays revealed a loss of cell death in *eds1*-2, *pad4*-1, and *adr1* mutants, whereas *sag101*-3 and *nrg1* mutants retained this response (fig. S15, E and F). The S_499_F amino acid exchange in the *pad4*-14 allele is in close proximity to the Rib-AMP/ADP binding surface of PAD4 (*40*) (fig. S15G) and might interfere with the formation of the EDS1-PAD4-ADR1 signaling hub. Analyses with an established pRIB-AMP/ADP binding mutant (PAD4_R314A_) (*40*) revealed that pRib-AMP/ADP-dependent EDS1-PAD4-ADR1 complex assembly is required for WTA-induced cell death formation (fig. S15H). This suggests that, similar to LRR-PRRs (*39*), CRKs also co-opted the EDS1-PAD4-ADR1 signaling node for defense signaling downstream of WTA perception.

## Discussion

Our results reveal that plants, akin to animals, are capable of specifically detecting Gram-positive bacteria via dedicated cell-surface receptors. In animals, recognition of lipoteichoic acid (LTA) is mediated by TLR2 (*7*), while wall teichoic acids (WTAs) are sensed by C-type lectin receptors, including LANGERIN, MANNOSE-BINDING LECTIN (MBL), and SURFACTANT PROTEIN D (SP-D) (*41*). We demonstrate that CRK7 is essential for WTA perception in the model plant *Arabidopsis thaliana*. CRKs form one of the largest RLK families, comprising 44 members in *Arabidopsis* and over 1,000 orthologs across diverse crop species (*42*). Their ectodomains harbor two DUF26 motifs with a conserved C-X_8_-C-X_2_-C signature (*43*), and certain DUF26-containing proteins have been reported to bind carbohydrates (*44, 45*). Identification of WTA as a Gram-positive–specific MAMP acting through a CRK receptor supports the hypothesis that CRKs can function as lectin-like carbohydrate-binding receptors in plant immunity (*42, 46*).

Gram-positive strains producing GroP-type WTAs induced more pronounced cell death than those with RboP-type WTAs. Furthermore, genetic evidence indicates that WTA glycosylation is required for cell death induction by *B. subtilis*, *S. aureus*, and *L. monocytogenes*. These findings suggest that, similar to human WTA receptors (*26*), CRK7 may discriminate specific glycosylation patterns to achieve binding specificity. We propose that plant-associated Gram-positive bacteria, including symbionts, commensals, and pathogens, may alter WTA composition and glycosylation to evade host immunity. In animal systems, WTA diversity is known to be shaped by selective pressures, including host adhesion and immune evasion, as structural variations modulate interactions with host receptors and immune components (*26, 41, 47*). Notably, in *S. aureus*, WTA also acts as a virulence factor by mediating adhesion to epithelial cells via host scavenger receptors, promoting biofilm formation and tissue colonization (*47*). An analogous strategy exploiting plant cell-surface receptors for adhesion or colonization has not yet been described but represents a plausible mechanism.

FLIM-FRET and co-immunoprecipitation analyses revealed that CRK7 forms homomers, resembling the multimeric human WTA receptor LANGERIN, where multimerization is essential for ligand binding (*48*). CRK7 also forms a preassembled receptor complex with the RLCK CRK43. We propose that ligand binding induces conformational changes in CRK7, triggering CRK43 dissociation. The model of a functional CRK7–CRK43 WTA receptor complex is supported by the observation that transgenic co-expression of both CRKs confers WTA responsiveness in *N. benthamiana*. Whether stable expression of this complex in solanaceous crops, such as tomato, can enhance resistance against Gram-positive pathogens like *C. michiganensis* (*11*) warrants further investigation.

Genetic data further indicate that CRK7-mediated WTA recognition depends on the EDS1–PAD4–ADR1 signaling hub, which is postulated to act as a convergence module that integrates pattern-triggered and effector-triggered immunity. The WTA insensitivity of the PAD4_R314A_ mutant suggests a requirement for small signaling molecules generated by TNL immune receptors (*40, 49*), though their origin remains unresolved. Our findings support a model (*39, 50*) in which specialized RLCKs, such as CRK43, short-circuit canonical immune signaling to promote defense priming and programmed cell death (Fig. 4G). This alternative pathway operates independently of Ca²⁺ influx, ROS production, and MAPK activation. Collectively, these results illustrate how plants flexibly and dynamically orchestrate immune responses in the absence of an adaptive immune system.

## Acknowledgments

We thank Jeff Erington (University of Sydney, Camperdown, NSW, Australia) for providing the *B*. *subtilis* 168 Δ*tagO* and Δ*ltaS* Δ*ynfl* Δ*yqgS* Δ*yvqJ* strains, Tom Grunert (University of Veterinary Medicine, Vienna, Austria) for providing the *S*. *aureus* wild type and mutant strains, Angelika Gründling (Imperial College London, London, UK) for providing the *L*. *monocytogenes* wild type and mutant strains, Slimane Choubane (Higher School of Biological Sciences of Oran, Oran, Algeria) for the *B*. *subtilis* TLO3 isolate, Ming Sun (Huazhong Agricultural University, Wuhan, Hubei, China) for the *B*. *subtilis* BSn5 isolate and Marcel Wiermer (Free University Berlin, Berlin, Germany) for providing the *Pst* DC3000 strain. We further thank Jane E. Parker (Max Planck Institute for Plant Breeding Research, Cologne, Germany) for providing the *pad4*-1/*sag101*-3 *P_PAD4_:YFP-PAD4_WT/R314A_* complementation lines and Till Ischebeck (University of Muenster, Muenster, Germany) for providing the tested *Arabidopsis* accessions. Lastly, we thank Fabian von Pappenheim (Göttingen University, Göttingen, Germany) and Jeroen Codée (Leiden University, Leiden, Netherlands) for discussions advancing the project, and lastly Robin Walz and Sabine Wolfarth for technical support. AI-assisted technology (ChatGPT-5) was used for text editing purposes.

## Funding

This work was supported by the Deutsche Forschungsgemeinschaft (DFG; German Research Foundation), grants RI 2920/3-1 to JR, GRK 2172 and INST 186/1277-1 FUGG (Confocal Microscope Leica TCS SP8) to VL. SK was supported within the frame of the joint international DFG-IRTG/NSERC-CREATE training program IRTG 2172 “PRoTECT: Plant Responses To Eliminate Critical Threats” (GRK 2172, project number 273134146) at the Göttingen Graduate Center of Neurosciences, Biophysics, and Molecular Biosciences (GGNB).

## Author contributions

Conceptualization: LP, JR, AM, CDC, EP, VL

Methodology: LP, AM, CDC, EP, VL

Investigation: LP, SK, CS, SA, LMS

Formal analysis: LP, SK, CS, SA, AM, CDC, EP, VL

Visualization: LP, VL

Funding acquisition: JR, VL

Project administration: VL

Supervision: VL

Writing – original draft: LP, VL

Writing – review & editing: LP, SK, CS, JR, AM, CDC, EP, VL

## Competing interests

Authors declare that they have no competing interests.

## Data and materials availability

All data are available in the main text or the supplementary materials.

## Supplementary Materials

### Materials and Methods

#### Plant growth conditions and plant material

Plants were grown on soil soaked once with 2ml/l Wuxal Super (Manna, Ammerbuch-Pfäffingen, Germany). *Arabidopsis* plants were grown under short-day conditions (8 h day period, 22 °C, 16 h night period, 18 °C) at 65% relative humidity. Other plant species used in this work were grown at long-day conditions (16 h day period, 25 °C, 6 h night period, 22 °C) at 65% relative humidity. T-DNA insertion lines of *Arabidopsis* were obtained from NASC. Mutant lines used in this work are stated in Table S1. *Arabidopsis* Ecotypes, listed in Table S2, were provided by Till Ischebeck (Westfälische Wilhelms-Universität-Münster, Germany).

#### Cloning

For complementation of the *crk7*-5 mutant with *pGreenII-0229-P_CRK7_:CRK7*, the genomic region of CRK7, including 1400 bp 5’ of the CRK7 start codon as a natural promoter region, and the full 3’UTR and terminator region were amplified from genomic DNA using primers oLP84 and oLP68. The *pGreenII-0229* vector was amplified using primers oLP83 and oLP85. Both amplified vector and insert were digested with restriction enzymes HindIII and AscI (Thermo Fisher) and ligated using T4 ligase (Thermo Fisher). For expression of mCitrine labeled CRK7, *mCitrine* was amplified using primers oLP159 and oLP160. The *pGreenII-0229-P_CRK7_:CRK7* plasmid was amplified using primers oLP161 and oLP162. Both PCR products were ligated using GIBSON assembly to generate *pGreenII-0229-P_CRK7_:mCitrine-CRK7*. For FLIM/FRET analysis, the mCitrine sequence of *pGreenII-0229-P_CRK7_:mCitrine-CRK7* was replaced by *mScarlet-I*. The *mScarlet-I* sequence (*51*) was amplified using primers oLP218 and oLP219. The *pGreenII-0229-P_CRK7_:mCitrine-CRK7* plasmid was amplified using primers oLP216 and oLP217. Both PCR products were assembled to generate *pGreenII-0229-P_CRK7_:mScarlet-I-CRK7* using GIBSON assembly. For complementation of the *crk43*-1 mutant with *pGreen-0229*-*P_CRK43_:CRK43_CDS_-mCitrine*, the CRK43 sequence was amplified from cDNA, using primers oSK37 and oSK38, with flanking EcoRI and NotI sites. The *pGreenII-0229* derivative containing *P_CRK43_* and mCitrine was digested with restriction enzyme EcoRI and NotI (Thermo Fisher), along with the *CRK43* PCR product. The digested products were ligated with T4 ligase enzyme (Thermo Fisher).

The plasmid used to generate the *B*. *subtilis* Δ*tagE* mutant straine was cloned by amplifying the upstream and downstream regions of *tagE* from genomic DNA using primers JR101 and JR102 or JR103 and JR104, respectively. The erythromycin (erm) resistance cassette was amplified from plasmid pDG646 (*52*) using primers mls-fwd and mls-rev. Long flanking region PCR was performed using primers JR101 and JR104 to fuse the upstream region, the erm cassette, and the downstream region.

#### Transient and stable plant transformation

All cloned plasmids were analyzed by sequencing before transformation into electrically competent *Agrobacterium tumefaciens* GV3101+pSoup (*53*). For transient expression in *N. benthamiana*, *A. tumefaciens* cultures were grown at 28°C to an optical density at 600 nm (OD_600_) between 0.8 and 1.2. Cells were harvested by centrifugation and resuspended to an OD_600_ of 0.6 in infiltration medium (10 mM MgCl_2_, 10 mM MES, 150 µM acetosyringone, pH 5.4). Resuspended cultures were incubated for 2 h at room temperature before syringe infiltration into the leaves of *N*. *benthamiana*. Infiltrated plants were kept at long-day conditions for 2-3 days or 5 days until further analyzed by microscopy and western blot, or macroscopic cell death evaluation, respectively.

Stable transformation of *Arabidopsis crk7*-5 and *crk43*-1 plants was performed using the floral dip method (*54*). Briefly, *A*. *tumefaciens* cultures were grown at 28 °C to an oD_600_ between 0.8 and 1.2. Cells were collected via centrifugation and resuspended in dipping solution (5% sucrose, 0.05% Silwet-L77). Inflorescent *Arabidopsis* plants were briefly dipped into the solution and kept at high humidity and low light for 16-24 h. Afterward, plants were grown in long-day conditions to harvest seed material. Transformants were screened using BASTA selection (*55*), Western Blot, and Confocal microscopy for mCitrine fusion proteins.

#### Transformation of *B*. *subtilis* 168

Transformation of *B*. *subtilis* strain 168 was performed as previously described (*56*). A culture of *B*. *subtilis* 168 was grown in MNGE medium (2% glucose, 0.2% potassium glutamate, 100 mM potassium phosphate buffer pH 7, 3.4 mM trisodium citrate, 3 mM MgSO_4_, 42 mM ferric ammonium citrate, 0.24 mM L-tryptophan, 0.1% casein hydrolysate) at 37 °C until the exponential growth phase shifted to stationary growth. The culture was diluted with an equal volume of MNGE medium lacking casein hydrolysate. After one additional hour of incubation at 37 °C, 2 µg of plasmid DNA (pBQ200 (*57*) empty vector) or PCR product (for generation of *B*. *subtilis* 168 Δ*tagE*) was added, and cells were incubated for 30 min at 37 °C. Expression solution containing 2.5% casein hydrolysate, 2.5% yeast extract, and 1.2 mM tryptophan was added. After 1 h of incubation, cells were transferred to lysogeny broth (LB) selection plates containing appropriate antibiotics.

#### Bacterial material, growth, and preparation

Gram-positive bacteria used for infiltration experiments were grown in LB medium at 37 °C to an OD_600_ of 0.5-2. For heat inactivation and infiltration, cells were harvested by centrifugation at 4000 g for 20 min, washed twice with 10 mM MgCl_2,_ and resuspended in 10 mM MgCl_2_ to an OD_600_ of 20. Resuspended cultures were heat-inactivated by autoclaving at 121 °C for 20 min. After heat inactivation, cultures were diluted to the respective OD_600_ mentioned in the figure legends and used for infiltration.

*Pseudomonas syringae* pv *tomato* DC3000 (*Pst* DC3000) was grown in NYG-medium containing 50 µg/ml rifampicin and 50 µg/ml kanamycin at 28 °C to an OD_600_ of 0.1-0.2. Cultures were harvested by centrifugation at 4000 g for 20 min, washed twice in 10 mM MgCl_2_, and resuspended in infiltration medium (10 mM MgCl_2_, 0.002% Silvet-L77) to 10^4^ for priming assays, or to 10^4^ or 10^7^ cfu/ml for comparison of phenotype induction of *B. subtilis* 168 pBQ200 empty vector. The *B. subtilis* 168 pBQ200 strain was used to allow the quantification of colony-forming units on erythromycin selection plates.

#### Fractionation of heat-inactivated *B. subtilis* cultures

Heat-inactivated *B. subtilis* cultures were precleaned by performing ultracentrifugation (Sorvall® WX Ultra Series; Thermo Fisher Scientific, Waltham, USA) at 100 000 g for 1 h. The supernatant was collected for infiltration into *Arabidopsis* leaves and further fractionation. Size-based fractionation was performed using Vivaspin® centrifugal concentrator columns (Sartorius Stedim, Goettingen, Germany) with a molecular cut of weight of 5 kDa or 30 kDa, according to the user’s manual. Flow-throughs were collected for infiltration into *Arabidopsis* Col-0 leaves. For the detection of mannose or glucose moieties, precleaned, heat-inactivated *B*. *subtilis* was incubated with 200 µl Concanavalin-A agarose beads (Cytiva, Marlborough, USA) and incubated at 4 °C for 16 h. After incubation, the supernatant was collected and used for infiltration. For proteinase treatment, pre-cleaned *B*. *subtilis* was mixed with 0.5 µg/µl BSA to allow visualization of proteinase activity via western blot. Samples were denatured at 95 °C for 20 min. Afterward, samples were treated with 0.05 µg/µl Proteinase K (New England Biolabs, Ipswich, USA), incubated at 45 °C for 4 h with a final inactivation of the Proteinase K at 95 °C for 20 min. The treated sample was used for SDS-PAGE analysis and infiltration. To determine the charge of the BSF, precleaned *B*. *subtilis* was run on an anion exchange column (Sartorius Stedim, Goettingen, Germany) according to the user’s manual. Flow-through was collected for infiltration. The column was washed with water and eluted with 5 M ammonium sulfate. The eluate was desalted using a PD10 gravity column (Cytiva, Marlborough, USA) according to the user’s manual before infiltration.

#### Size exclusion chromatography-based WTA purification

WTA was purified from heat-inactivated *B*. *subtilis* pre-cleaned via ultracentrifugation through a combination of chromatographic techniques. Size exclusion chromatography was performed on Sephacryl® S-300 HR resin (1.5 x 45cm, Cytiva 17-0599-01), packed and eluted with ammonium bicarbonate 50mM. The flow rate was set to 15 ml/h, and the eluate was monitored by a refractive index detector. A subsequent anion exchange chromatography on Q-sepharose® Fast-Flow (1 x 1.5 cm, Cytiva 17-0510-01) was useful to isolate the teichoic acids; the elution was achieved by a stepwise gradient of NaCl (10, 100, 200, 400, 700, 1000 mM). Each solution was concentrated by freeze-drying and desalted on BioGel® P-10, run in water with a flow rate of 12 ml/h, and the eluate was monitored by a refractive index detector.

#### NMR analysis

^1^H and 2D NMR spectra were recorded using a Bruker 600 MHz spectrometer equipped with an inverse cryoprobe with gradients along the z-axis. The samples were solved in 550 µl of D_2_O, and the spectra were calibrated with internal acetone (dH= 2.225 ppm; dC = 31.45 ppm). Total Correlation Spectroscopy (TOCSY) and Nuclear Overhauser Enhancement Spectroscopy (NOESY) experiments were performed using data sets (t1 × t2) of 2048 × 512 points. Heteronuclear Single-Quantum Coherence (HSQC) and Heteronuclear Multiple Bond Correlation (HMBC) experiments were performed in the ^1^H-detection mode by single-quantum coherence with proton decoupling in the ^13^C domain using data sets of 2048 × 512 points. HSQC was performed using sensitivity improvement and the phase-sensitive mode using echo/antiecho gradient selection, with multiplicity editing during the selection step (*58*). HMBC was optimized on long-range coupling constants, with a low-pass J filter to suppress one-bond correlations, using gradient pulses for selection, and a 60ms delay was used for the evolution of long-range correlations. HMBC spectra were optimized for 6 – 15 Hz coupling constants. For transformation, the data matrix in both homo-and heteronuclear experiments was extended to 4096 × 2048 points and transformed by applying a qsine or a sine window function (*59*). Spectra were recorded at 298 K. TOCSY and T-ROESY experiments were performed with a mixing time of 100 and 400 ms, respectively. 40 scans for COSY, 32 for TOCSY, 64 for HSQC, and 80 for HMBC were recorded. All the spectra were transformed and analyzed with Bruker Topspin NMR DATA analysis software.

#### Large-scale expression of WTA from *B*. *subtilis*

High-quantity extraction of WTA from *B*. *subtilis* was performed as previously described (*60*), with mild modifications. 5 L of *B*. *subtilis* culture was harvested by centrifugation, washed with 1 M NaCl, and resuspended in 1 M NaCl. The re-suspended bacteria were disrupted using a French Press cell disruptor. Cell fragments from the disrupted culture were collected by centrifugation. Cell fragments were recovered in water and stirred at 60 °C for 30 min to remove non-covalently bound compounds. Proteins were removed via incubation in 0.2 mg/ml trypsin at 37 °C for 18 h. To hydrolyze WTAs from the peptidoglycan cell wall, the sample was incubated in 10% w/v trichloroacetic acid at 4 °C for 18 h. Peptidoglycan was removed by centrifugation, and WTAs were precipitated from the supernatant with 0.1 volumes of 3 M sodium acetate, pH 5.2, and 3 volumes of 95% ethanol. The mixture was incubated overnight at −80 °C and centrifuged to harvest precipitated WTAs. The pellet was washed with 95% ethanol and re-suspended in water. Solved WTAs were transferred into a fresh tube and lyophilized to determine the yield quantity.

#### Ion leakage assay

One leaf per plant was infiltrated with the bacterial strain of interest or WTA (12 leaves per plant line and treatment), and the plants were grown for 5 days under short-day conditions. Leaves were harvested into 2 ml reaction tubes containing 1.5 ml deionized water and incubated for an additional 16 h, gently shaking at room temperature. Conductivity (C_sample_) was measured using a Seven-2 Go conductivity meter S7 equipped with an InLab 751-4 mm platinum probe (Mettler-Toledo GmbH, Gießen, Germany). After the first measurement, each sample was autoclaved at 121 °C for 20 minutes, subsequently cooled to room temperature, and the maximum conductance (C_max_) was measured. For each measurement, normalized conductivity was calculated by dividing C_sample_ by C_max_.

#### Arabidopsis accession screen

Three individual plants of each Arabidopsis accession were grown under short-day conditions. After 5 weeks, 3 leaves of each individual plant were infiltrated with pre-purified *B. subtilis* 168 adjusted to an OD_600_ of 0.02 in 10 mM MgCl_2_. Sensitive and insensitive accessions were determined by visual examination at 6 dpi. Insensitive accessions were regrown for ion-leakage measurements.

#### ROS burst assay

To measure ROS accumulation after PAMP treatment, leaf discs (4 mm diameter) of Col-0 plants were transferred into wells of a 96-well plate filled with water. The plates were incubated overnight. The next day, the water was replaced with a luminol solution (50 µM L-012 and 10 µg/ml horse radish peroxidase) containing 100 nM flg22, 1µg/ml WTA, or water as a control. Luminescence was measured using a TECAN Infinite® M200 plate reader (Tecan Group Ltd, Maennedorf, Switzerland).

#### Cytoplasmic Ca^2+^ assay

Cytoplasmic Ca^2+^ measurements were performed as previously described (*61*). Seedlings of *Arabidopsis* Col-0 plants expressing aequorin were grown in vitro on semi-solid ½ MS + sucrose agar plates. Seven to nine days old seedlings were transferred to a 96-well plate with 100 µl of 20 µM coelenterazine dissolved in water and incubated in the dark for 16 h. After incubation, the coelenterazine solution was replaced with water, and the seedlings were incubated in the dark for an additional 30 minutes. Seedlings were treated with 1 µg/ml WTA, 100 nM flg22, or water as a control. Luminescence (L) was measured using a TECAN Infinite® M200 plate reader (Tecan Group Ltd, Maennedorf, Switzerland). Maximum luminescence (L_max_) was assessed after treatment with discharge solution (10% Ethanol, 1 M CaCl_2_).

#### RNA extraction and RT PCR

For qRT-PCR, leaves were treated with 1 µg/ml WTA or 10 mM MgCl_2_. Three infiltrated leaves of different plants were harvested as one biological replicate two days after infiltration. Leaf material was ground, and RNA extraction was performed using the InnuPREP Plant RNA Kit (Analytik Jena AG, Jena, Germany), according to the user’s manual. Synthesis of cDNA was performed using RevertAid H Minus M-MuLV Reverse Transcriptase (Thermo Fisher Scientific, Schwerte, Germany). qRT-PCR was performed as described (*62*). PR1 and UBQ5 were amplified using previously described primers (*63*). Calculations were performed using the 2^−ΔΔCT^ method (*64*). The UBQ5 transcript level was used as a reference.

#### Sequencing of the *crk43*-1 splice site mutation

To identify the frame shift resulting from the crk43-1 splice site mutation, cDNA was prepared as described for RT PCR. The cDNAs of *CRK43* and *crk43*-1 were amplified using primers oCT87 and oCT96. The splice site mutation was sequenced using primer oCT96. Sequencing results were mapped to the wild-type CRK43 sequence using the Geneious software.

#### Priming assay

Priming assay was performed in *Arabidopsis* plants by pretreating leaves with 10 ng/ml WTA, 100 nM flg22, or Mock treatment 24 h before infiltration with *Pst* DC3000 at a final cell density of 10^4^ cfu/ml, as previously described (*65*). Colony-forming units per leaf area (cfu/cm^2^) were quantified by harvesting 8mm leaf discs from three infiltrated leaves from different plants as one sample at 3 dpi with *Pst* DC3000 and plating a dilution series on NYG plates containing 50 µg/ml rifampicin and 50 µg/ml kanamycin.

#### Confocal microscopy and FLIM measurements

Confocal laser scanning microscopy was performed as described (*51*) using a Leica Microsystems TCS SP8 Falcon system with LASX 3.3.3 software, equipped with a Leica HC PL APO x40/1.10 W CORR CS2 objective. Fluorophores were excited using a pulsed white light laser at 514 nm for mCitrine and 561 nm for mScarlet-I. Fluorophores were detected via HyD SMD detectors with a detection window of 525-555 nm or 580-660 nm for mCitrine and mScarlet-I, respectively. Images were taken in sequential scanning mode to avoid bleach-through of mCitrine signal into the mScarlet-I detection window. As expression of mCitrine-tagged CRK7 in Arabidopsis is barely detectable via microscopy, Arabidopsis plants used for the analysis of CRK7 localization were treated with *B. subtilis* at an OD_600_ of 0.02 two days before microscopy to induce CRK7 expression. Fluorescence Lifetime Imaging (FLIM) was measured as described (*51*). The same lasers and detectors were used as for confocal imaging. FLIM measurements were performed using the LASX single-molecule detection module (Leica). Protein binding was calculated from FRET analysis as previously described (*51*).

#### Western blot and co-immunoprecipitation

Protein extraction and western blot experiments were conducted as previously described (*66*). Primary antibodies used were rat monoclonal anti-GFP (Chromotek, 3H9) for the detection of mCitrine-tagged CRK7 and CRK43, rat monoclonal anti-RFP (Chromotek, 5F8) for the detection of mScarlet-tagged CRK7, and rabbit polyclonal anti-CRK7 (Agrisera) for the detection of untagged CRK7. Goat anti-rat AP-conjugate (Sigma-Aldrich, A8438) or goat anti-rabbit AP-conjugate (Sigma-Aldrich, A3687) were employed as secondary antibodies. The Immunstar-AP substrate (Bio-Rad, 1705018) was used to visualize signals on a Bio-Rad Chemidoc system. GFP pull-down experiments were performed as described by (*51*). Standard protein extracts were mixed with GFP-Trap agarose beads (Chromotek, gta). Beads were incubated overnight. After several washing steps, proteins were eluted in 1X SDS loading dye directly prior to gel loading.

#### Bioinformatic and Statistical analysis

All experiments were carried out at least 3 times, with the exception of western blot and microscopy of mCitrine tagged CRK7 complementation lines (fig. S9, B and C) as well as plant species (fig. S11) and Arabidopsis accession (fig. S12) infiltrations, which were carried out once. All experimental repeats yielded similar results. Statistical analyses of data and graph generation were conducted using the GraphPad Prism 8 software (GraphPad Software Inc.). Mapping-by-sequencing was performed by mapping 150bp paired-end or 50bp non-paired reads to the *Arabidopsis* reference genome using the CLC genomic workbench 10.1.1 (Qiagen, Hildern, Germany).

### Supplementary Text

#### 2D NMR structural studies of *B*. *subtilis* wall teichoic acid purification

The structure of the wall teichoic acids was determined by analyzing ^1^H, ^1^H homo- and ^1^H, ^13^C heteronuclear 2D NMR experiments, recorded by dissolving glycans in D_2_O. ^1^H,^1^H COSY, and ^1^H, ^1^H TOCSY experiments were used to disclose the protons of each spin system, while carbon atoms were identified through ^1^H, ^13^C HSQC. The HSQC spectrum presented one anomeric signal at 5.19 ppm, labeled with the capital letter A (Fig. S5b). In the TOCSY spectrum (Fig. S5d), correlations were present, with that at 3.55 ppm in common with the COSY spectrum. Hence, this cross peak was assigned to H-2, and by a similar approach, H-3 (3.78 ppm), H-4 (3.42 ppm), H-5 (3.95), and H-6,6’ (3.90-3.78). This TOCSY pattern is typical of a gluco-configured sugar, and this information, along with the chemical shift values from literature (*59*), led to identifying A as a terminal α-glucose. Finally, the study of the HMBC spectrum allowed the identification of residues B, C, D, and E (Fig. S5c) as, respectively, a free-linked 1,3-polyglycerol phosphate, a 2-linked 1,3-glycerol phosphate, a 2-linked 1-phosphate terminal glycerol, and a free 1-phosphate terminal glycerol. Chemical shift values of these residues (Tab. S5) matched with those reported in literature (*67, 68*). Since the intensities of the signals relating to residues D and E were lower than those of the signals of B and C, they could represent chain terminators or fragments of chains hydrolyzed during the various work-up procedures. In conclusion, the main signals of the NMR spectra belong to the poly-gro-type polymer. The backbone of this polymer is made of 1,3-linked polyglycerolphosphate (units B and C); the position 2 of some of these Gro units is further substituted with an α-Glucose (unit A). By integration of the B2 vs. C2 densities in the HSQC spectrum (Fig. S5b), it has been possible to evaluate the free-Gro/bound-Gro ratio, which is 36%.

---

**Fig S1.**
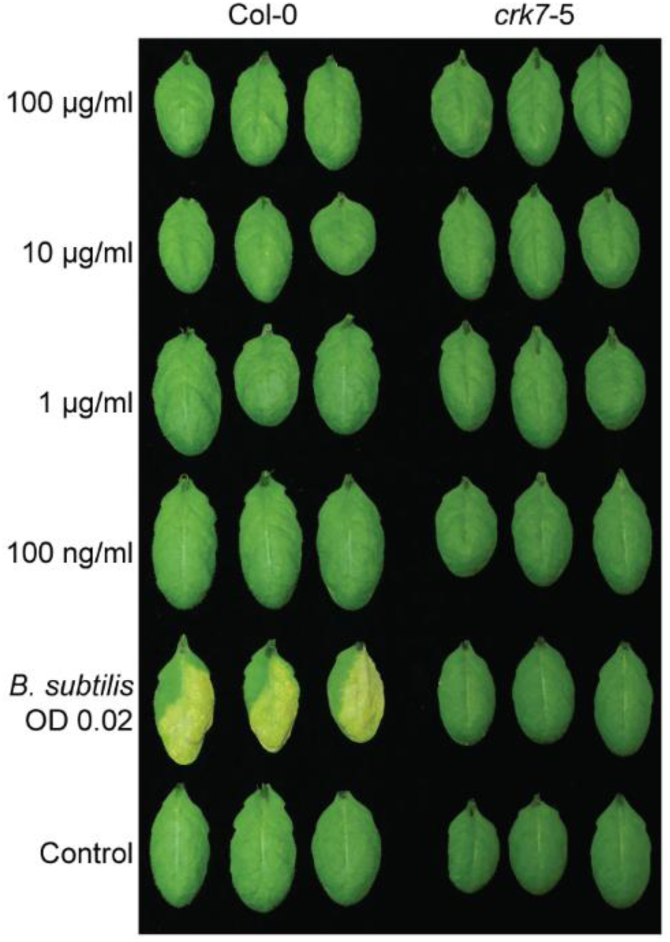
Surfactin infiltration does not induce cell death in *Arabidopsis*. Leaves of *Arabidopsis* Col-0 and the *crk7*-5 mutant were infiltrated with Surfactin at concentrations from 100 µg/ml to 100 ng/µl. Heat-inactivated *B*. *subtilis* at an OD_600_ of 0.02 in 10 mM MgCl_2_ and 10 mM MgCl_2_ were infiltrated as positive and negative controls, respectively. Pictures show three infiltrated leaves from three individual plants at 6 dpi.

**Fig S2.**
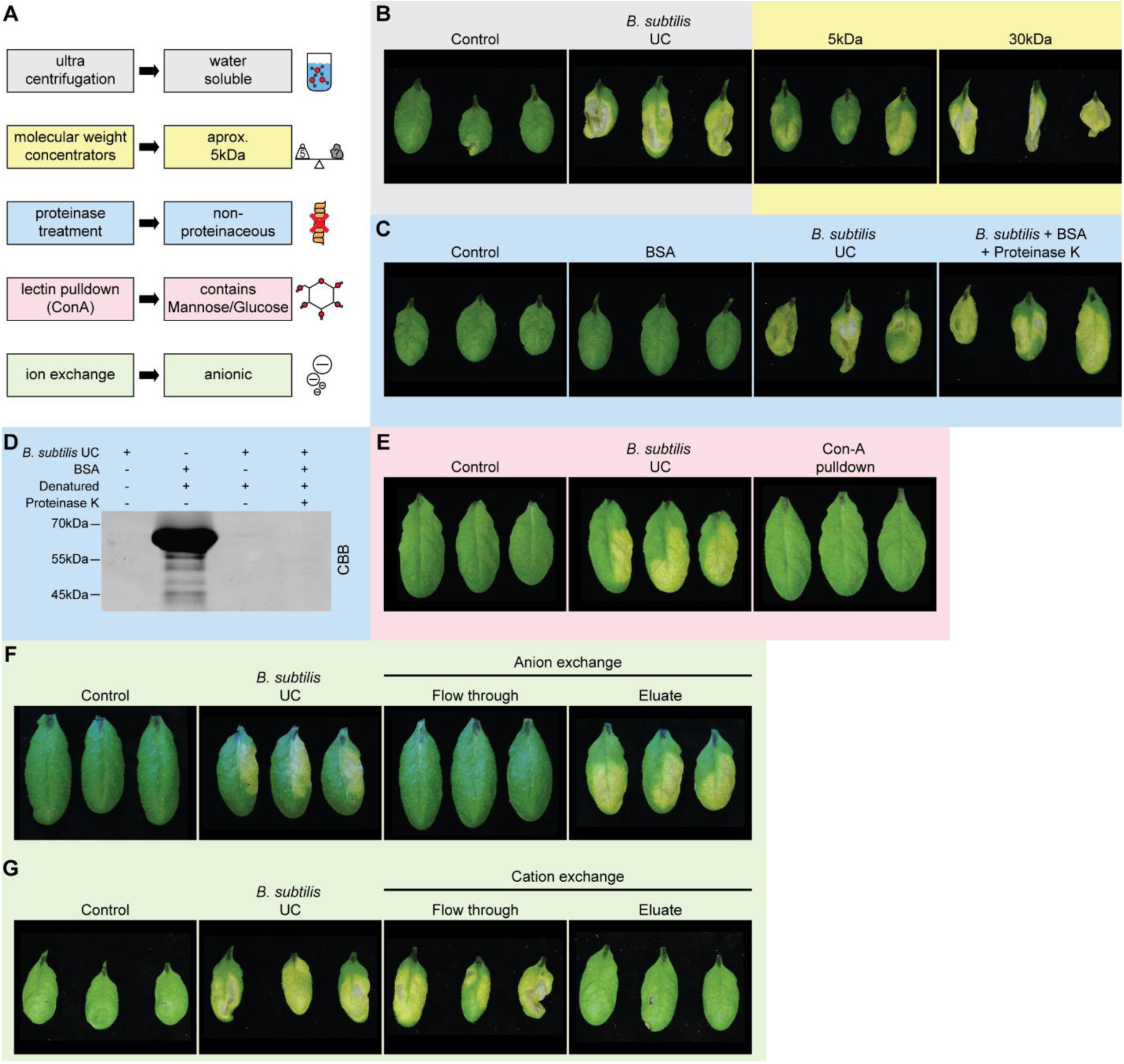
Biochemical analysis suggests the carbohydrate nature of the cell death inducing agent. (**A**) Scheme summarizing biochemical analysis of autoclaved *B. subtilis* cultures. (**B**) *B. subtilis* culture at an OD_600_ of 0.02 in 10 mM MgCl_2_, precleaned by ultracentrifugation (*B. subtilis* UC), was run through protein concentrator columns with a molecular weight cut-off at 5 kDa or 30 kDa. Flow through was used for infiltration. (**C** and **D**) *B. subtilis* UC was treated with Proteinase K. BSA was added as a control for Proteinase K activity. (C) Cell death-inducing capacity was evaluated by infiltration, and (D) Proteinase K activity was monitored via SDS-PAGE stained with Coomassie Brilliant Blue (CBB). (**E**) *B. subtilis* UC was incubated overnight with Concanavalin A (Con-A) beads. The supernatant was used for infiltration. (**F** and **G**) *B. subtilis* UC was run through an anion- (F) and cation- (G) exchange column. Flow-through and desalted eluate (eluted with 5 M NH_4_SO_4_) were used for infiltration. (B, C, E, F, G) *B. subtilis* UC, or 10 mM MgCl_2_ were used as positive and negative controls, respectively. Images show three infiltrated leaves of three individual plants at 6 dpi.

**Fig. S3.**
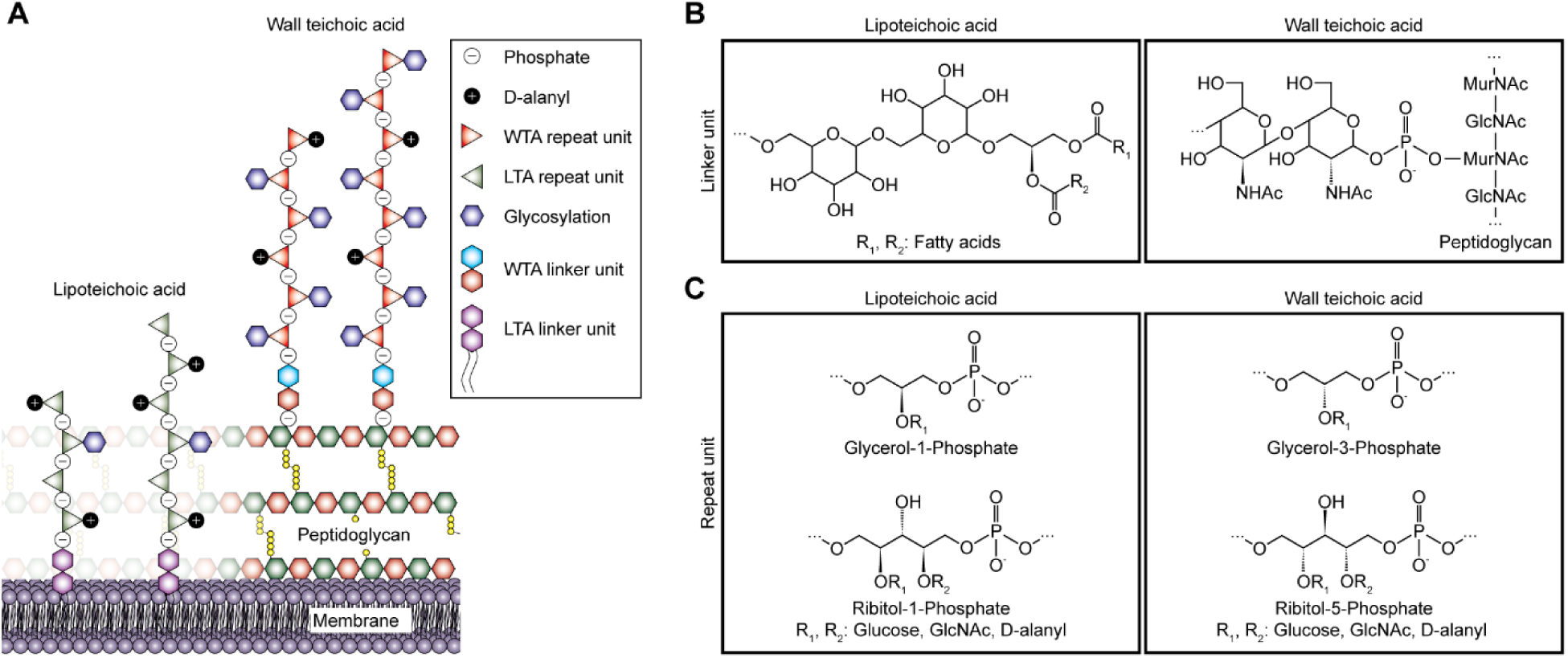
Schematic depiction of lipo- and wall teichoic acids. (**A**) Illustration of LTA and WTA at the cell envelope of Gram-positive bacteria. (**B**) Structural representation of the commonly characterized LTA- and WTA-linker units, consisting of glucosyl-β-1,6-glucosyl-β-1,3-diacylglycerol (*6*) and *N*-acetylmannosamine-β-1,4-N-acetylglucosamine-1-phosphate (*16*), respectively. (**C**) Chemical structure of the most common LTA and WTA repeat units, which are built by 1,3-poly-(glycerol-1-phosphate) or 1,5-poly-(ribitol-1-phosphate) and 1,3-poly-(glycerol-3-phosphate) or 1,5-poly-(ribitol-5-phosphate), respectively (*6, 16, 22*). Both LTA and WTA repeat units can be decorated by D-alanylation or varying glycosylation, including glucose or N-acetylglucosamine moieties (*6, 16*).

**Fig. S4.**
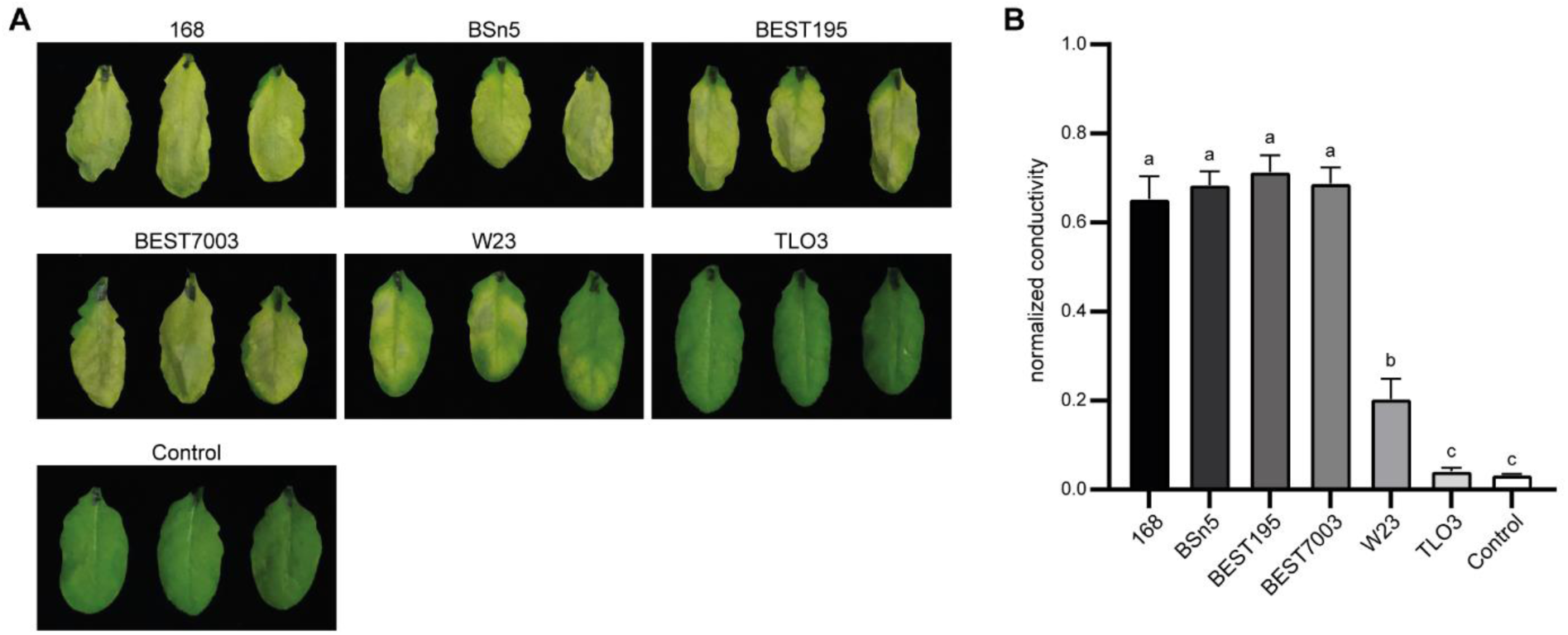
A *Bacillus subtilis* isolate lacking the WTA biosynthesis gene cluster does not induce cell death. (**A** and **B**) Infiltration of *Arabidopsis* Col-0 leaves with different *B*. *subtilis* isolates predicted to produce glycerol-phosphate-based (168, BSn5, BEST195, BEST7003), ribitol-phosphate-based (W23), or no wall teichoic acid (TLO3), according to the analysis of the WTA biosynthesis gene cluster (*18*). Each isolate was infiltrated at an OD_600_ of 0.02 in 10 mM MgCl_2_. 10 mM MgCl_2_ was infiltrated as a control. (A) Pictures show three infiltrated leaves from three individual plants at 6 dpi. (B) Ion leakage assay to quantify cell death at 6 dpi. Bars represent mean normalized conductivity ±SEM (n=12). Different letters indicate significant differences of means (One-way ANOVA with Tukey’s post hoc test, p<0.05).

**Fig. S5.**
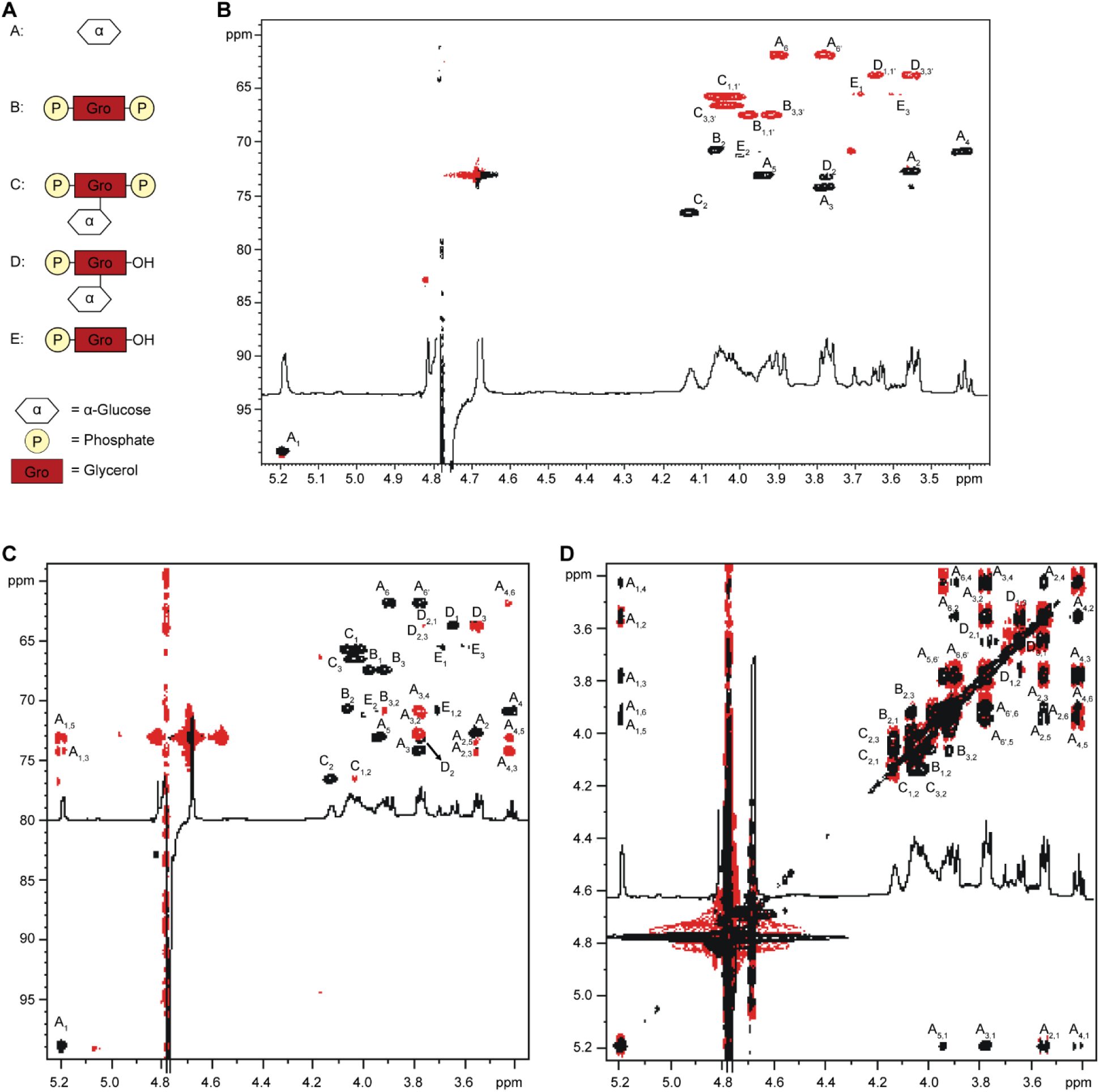
NMR analysis of WTAs purified from pre-cleaned *B*. *subtilis*. (**A**) Structures of carbohydrate residues identified by NMR analysis. (**B**) HSQC spectrum, along with the NMR profile. Red densities represent the –CH_2_ groups. (**C**) ^1^H, ^13^C HMBC spectrum in red and ^1^H, ^13^C HSQC spectrum in black. (**D**) ^1^H, ^1^H COSY spectrum in red and ^1^H, ^1^H TOCSY spectrum in black. (B to D) (600 MHz, 298 K, D_2_O) Letters refer to the carbohydrate residues reported in (A), and arabic numerals refer to the proton/carbon atoms of the respective residue. Signals marked with (*) are due to minor impurities.

**Fig. S6.**
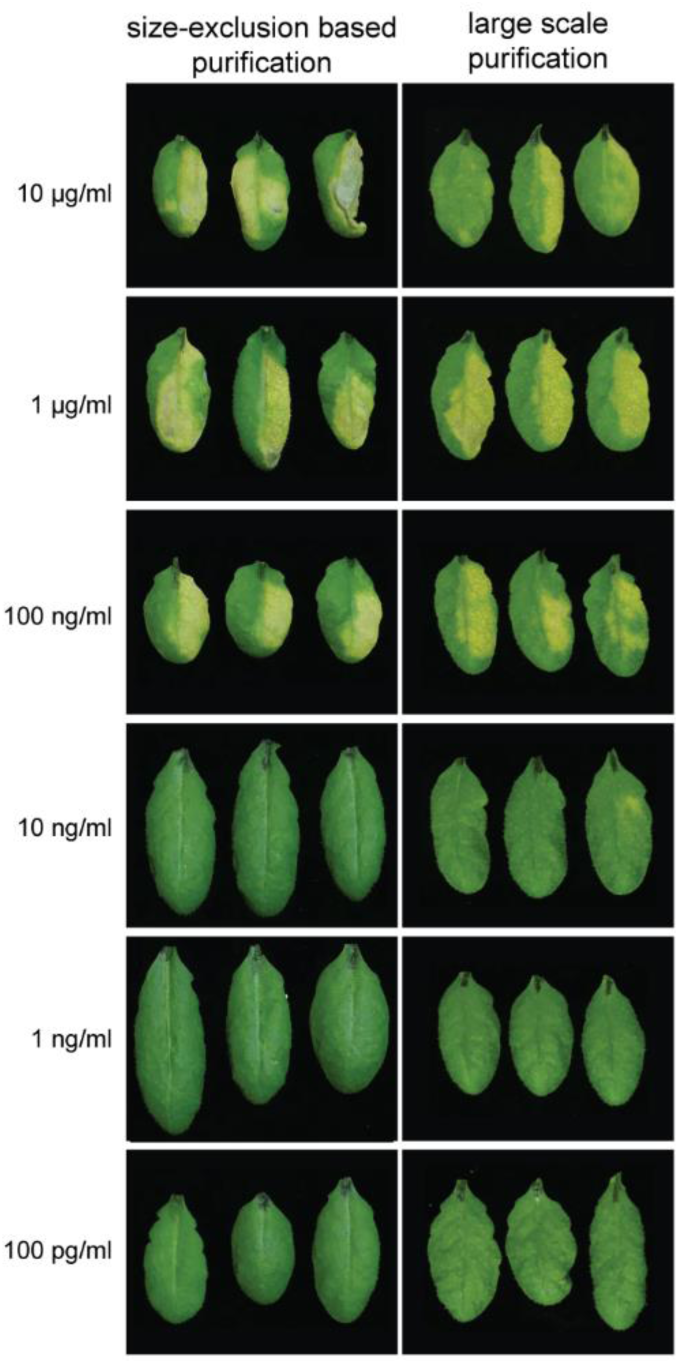
Large-scale WTA preparation yields WTA with cell death inducing activity. Infiltration of *Arabidopsis* Col-0 plants with WTA prepared via size exclusion chromatography (fig. S4) or by large-scale extraction from *B*. *subtilis* 168. Purified WTAs were infiltrated at concentrations from 10 µg/ml to 100 pg/ml. Pictures show 3 infiltrated leaves of 3 individual plants at 6 dpi.

**Fig. S7.**
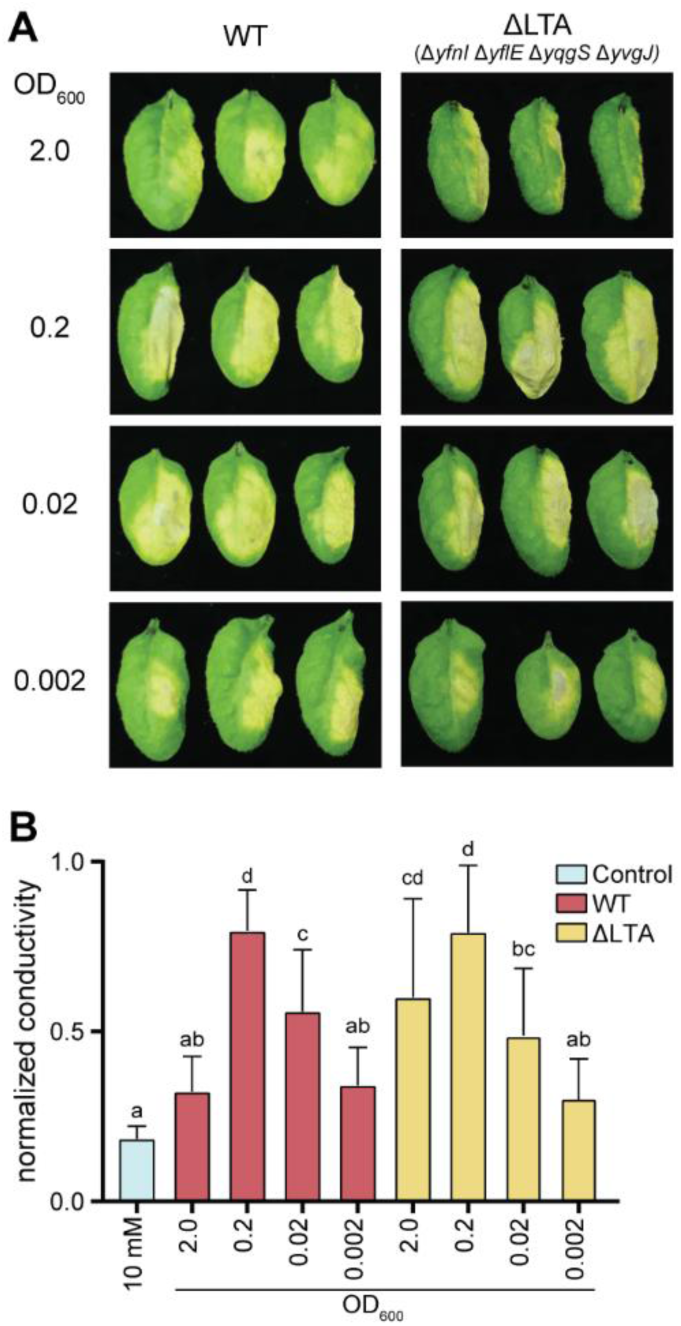
A *Bacillus subtilis* lipoteichoic acid biosynthesis mutant shows wild-type-like cell death induction in *Arabidopsis*. (**A** and **B**) Infiltration of *Arabidopsis* Col-0 with *Bacillus subtilis* 168 wild type and the lipoteichoic acid biosynthesis mutant Δ*ltaS* Δ*ynfl* Δ*yqgS* Δ*yvqJ* at an OD_600_ of 2.0 to 0.002 in 10 mM MgCl_2_. 10 mM MgCl_2_ was used as a control. (A) Images show three infiltrated leaves of three individual plants at 6 dpi. Only one-half of each leaf was infiltrated. (B) Ion leakage assay to quantify cell death at 6 dpi. Bars represent mean normalized conductivity ±SEM (n=12). Different letters indicate significant differences of means (One-way ANOVA with Tukey’s post hoc test, p<0.05).

**Fig. S8.**
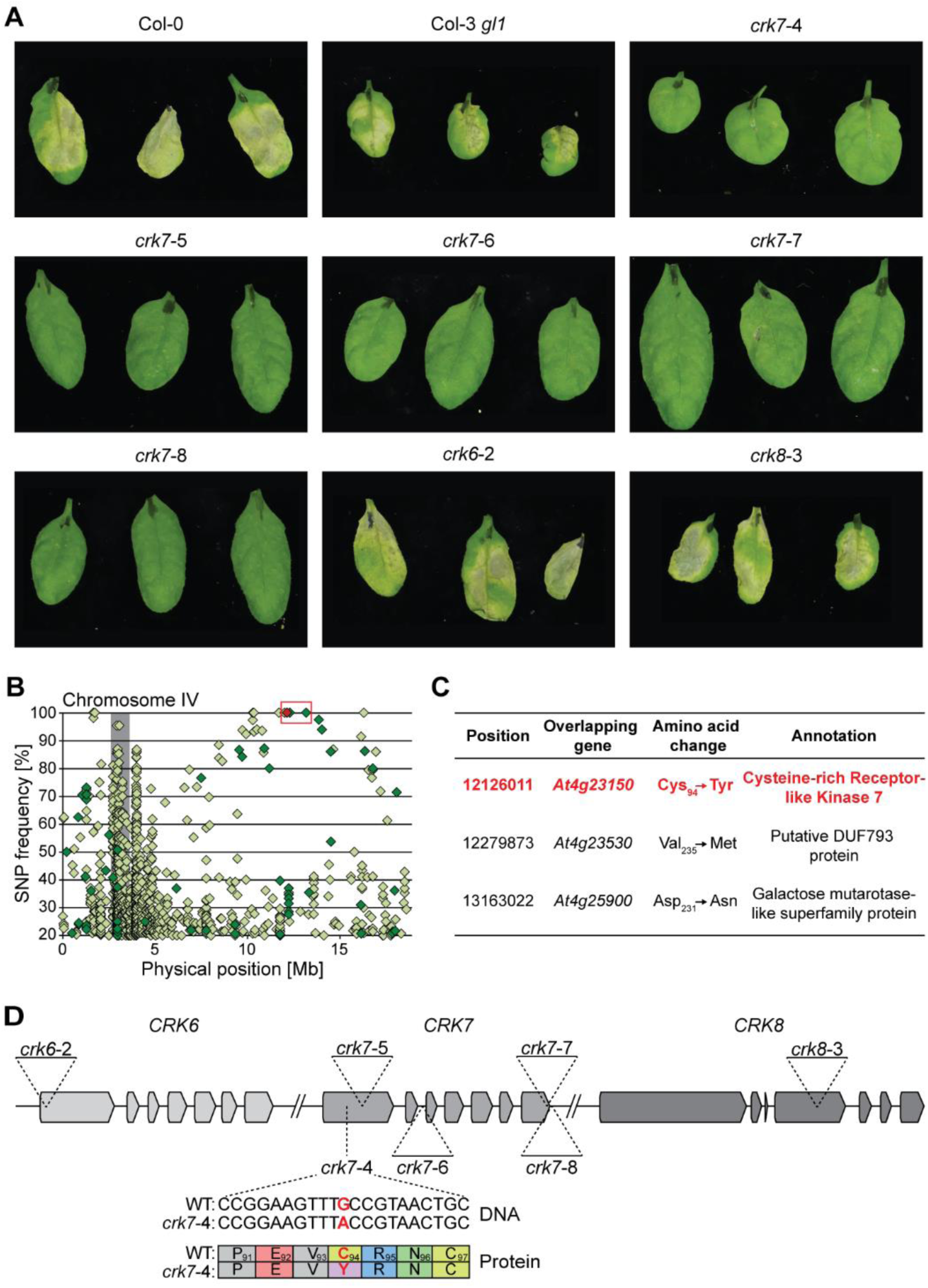
CRK7, but not CRK6 and CRK8, is required for WTA-induced cell death. (**A**) Infiltration of indicated mutants and corresponding wild-types (Col-0 for T-DNA lines, Col-3 *gl1* for *crk7*-4) with 1 µg/ml WTA in 10 mM MgCl_2_. Pictures show three infiltrated leaves of three individual plants at 6 dpi. (**B**) Identification of the *crk7*-4 mutant using mapping by sequencing of F2 plants with a suppressor phenotype. Diamonds indicate the position and frequency of synonymous (light-green) and nonsynonymous (dark-green) single-nucleotide polymorphisms (SNPs). Candidate SNPs are highlighted by the red box. The red diamond indicates the *crk7*-4 mutation. Grey background indicates the centromere region. (**C**) Candidate suppressor nsSNPs identified on Chromosome IV via mapping-by-sequencing. (**D**) Schematic representation of the *CRK6*-*CRK7*-*CRK8* gene cluster on Chromosome 4 of *Arabidopsis thaliana* Col-0. T-DNA insertion loci of *crk6*-2, *crk7*-5, *crk7*-6, *crk7*-7, *crk7*-8, and *crk8*-3 are depicted, as well as the C_94_Y amino acid exchange in the *crk7*-4 mutant.

**Fig. S9.**
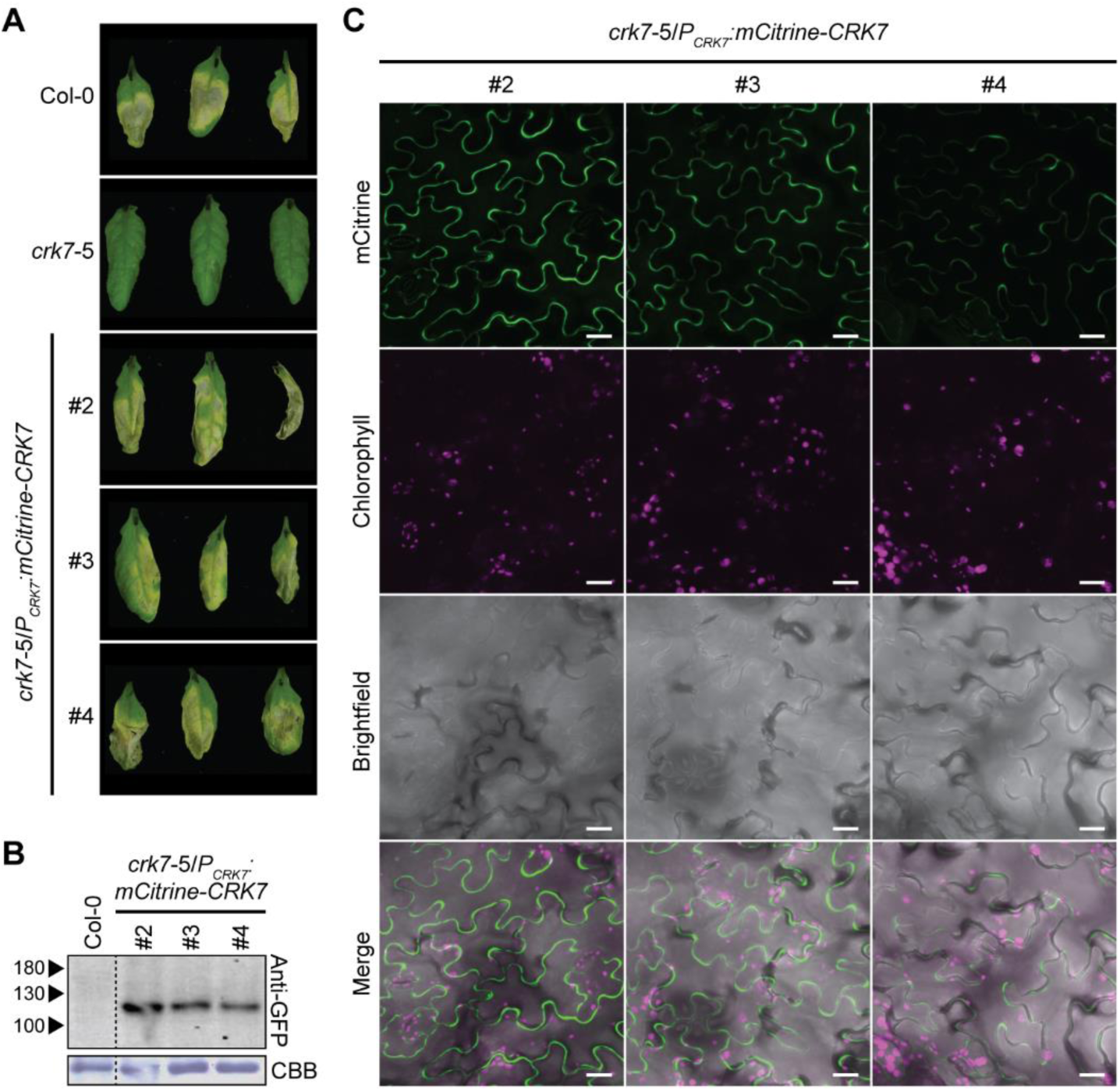
mCitrine-CRK7 localizes to the plasma membrane and complements the *crk7*-5 mutant phenotype. (**A**) Infiltration of three independent *crk7*-5 lines expressing mCitrine-tagged CRK7 under the control of the native promoter (*crk7*-5/*P_CRK7_:mCitrine-CRK7*). Col-0 and the *crk7*-5 mutant were infiltrated as controls. Plants were infiltrated with 1 µg/ml WTA in 10 mM MgCl_2_. Pictures show three infiltrated leaves of individual T3 plants at 6 dpi. (**B**) Western Blot analysis with anti-GFP antibody to confirm mCitrine-CRK7 expression in the tested complementation lines. (**C**) Confocal laser scanning microscopy of the three independent *crk7*-5/*P_CRK7_:mCitrine-CRK7* lines. mCitrine was excited at 514 nm and detected at 525 nm to 555 nm. Chlorophyll autofluorescence was detected at 740 nm to 770 nm. Fluorescence images represent maximum projections of 12 focal planes at 1 µm distance. Scale Bar = 20 µm.

**Fig. S10.**
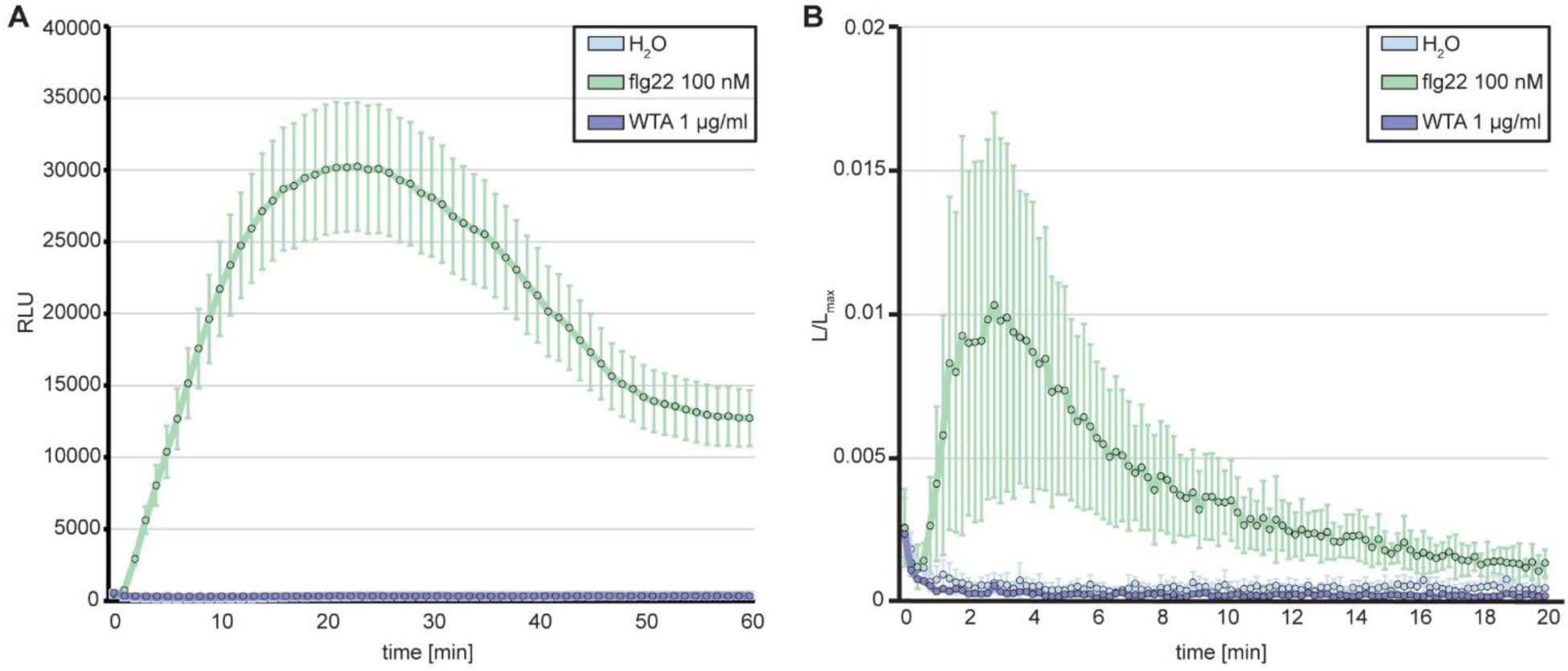
WTA treatment does not induce ROS burst and Ca^2+^ influx. Measurement of ROS production (**A**) and cytoplasmic calcium (**B**) after treatment of leaf discs from 5-week-old *Arabidopsis* Col-0 plants (A) or 7-9-day-old seedlings of Col-0 plants expressing aequorin (B) with 1 µg/ml WTA. (A and B) 100 nM flg22 and water treatments were used as positive and negative controls, respectively. (A) Data represent average relative light units (RLU) ± SEM (n=8). (B) Data are represented as average L/Lmax-normalized values ±SD (n=8).

**Fig. S11.**
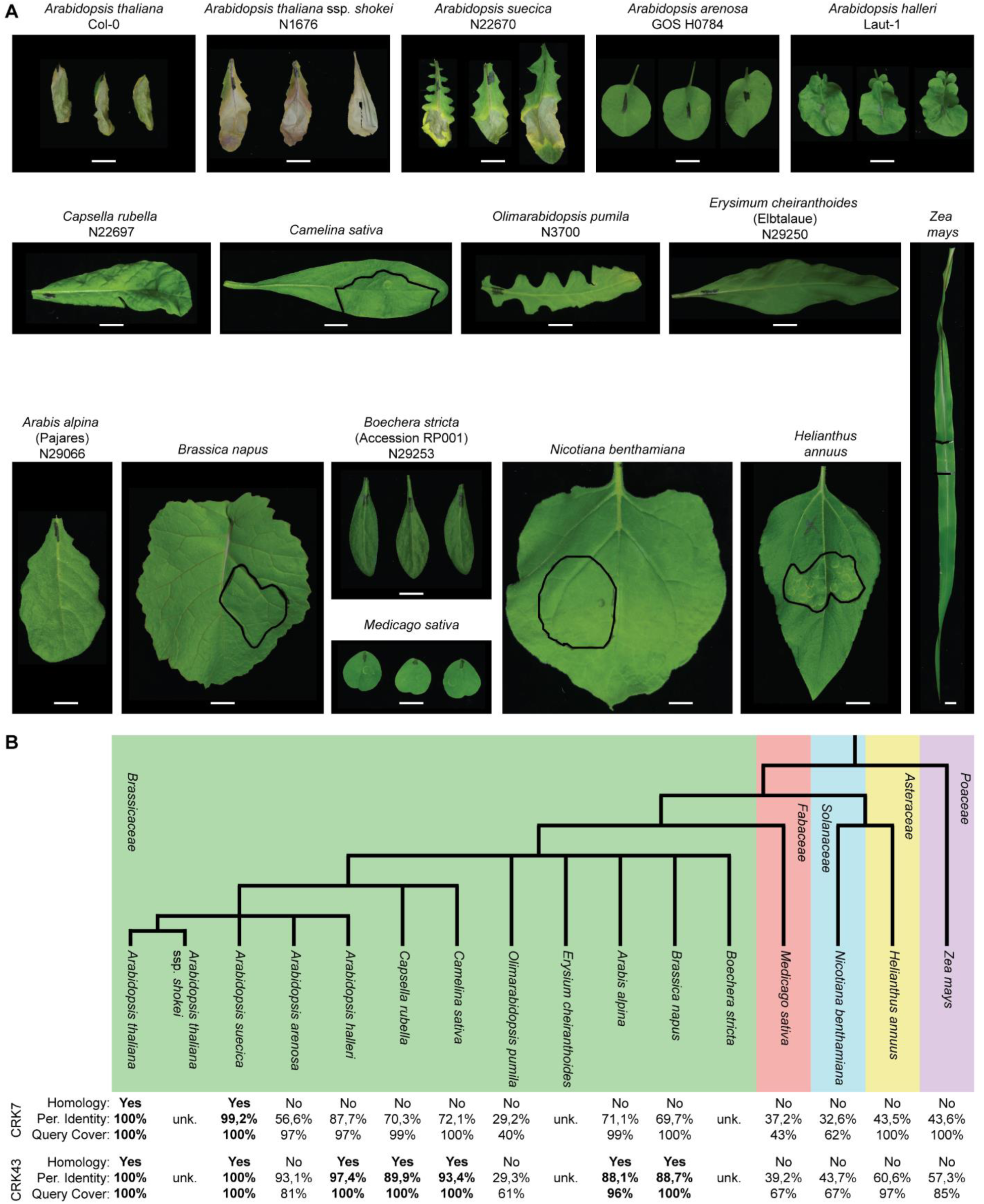
WTA-induced cell death is specific for *Arabidopsis* species, with a CRK7 homolog. (**A**) Different plant species were infiltrated with 1 µg/ml WTA in 10 mM MgCl_2_. Pictures show infiltrated leaves at 10 dpi. Black circles indicate the infiltrated area. If no circle is present, the whole leaf was infiltrated. (**B**) Taxonomic relationship of infiltrated plant species. The taxonomy tree was built using the NCBI Taxonomy Browser. BLASTp search was conducted using CRK7 and CRK43 protein sequences to identify CRK7 and CRK43 homologs in the analyzed species. Homology was accepted for hits with at least 85% identity, 95% query coverage, and a max-score per amino acid of at least 1.75.

**Fig. S12.**
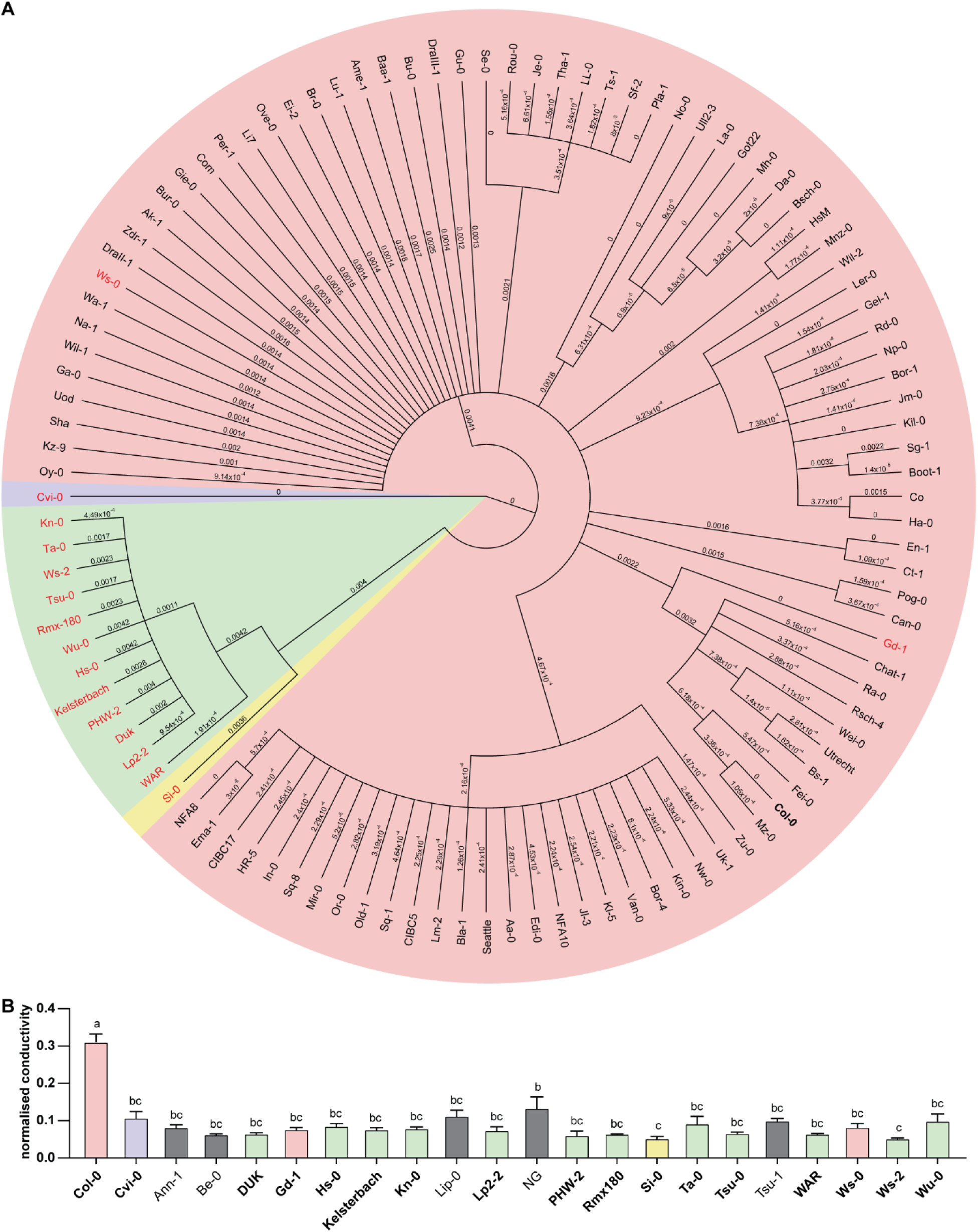
WTA-insensitive *Arabidopsis* accessions share homology in the CRK7 gene sequence. (**A**) Phylogenetic tree of the *CRK7* (*At4g23150*) genomic sequence of screened WTA-sensitive (black font) and insensitive (red font) *Arabidopsis thaliana* accessions for which a genome sequence was available from the 1001 genomes project (*29*). Neighbor joining tree was built using the Jukes-Cantor model based on a MUSCLE alignment of the full-length genomic region of CRK7, performed in Geneious©. The *At4g23150* locus of the Cvi-0 accessions was set as an outgroup; branch labels indicate substitutions per site. The CRK7 gene of different accessions was separated into four groups with no CRK7 gene present (blue background), functional CRK7 (red background), potentially non-functional group I (green background), and II (yellow background). (**B**) Ion leakage assay of WTA-insensitive *Arabidopsis thaliana* accessions. Col-0 was included as a sensitive accession control. Accessions included in the phylogenetic tree are labeled in bold letters. Bar color represents the corresponding group as depicted in (A). Leaves were infiltrated with heat-inactivated *B*. *subtilis* at an OD_600_ of 0.02 in 10 mM MgCl_2_. Bars represent mean normalized conductivity ± SEM (n=12). Different letters indicate significant differences of means (One-way ANOVA with Tukey’s post hoc test, p<0.05).

**Fig. S13.**
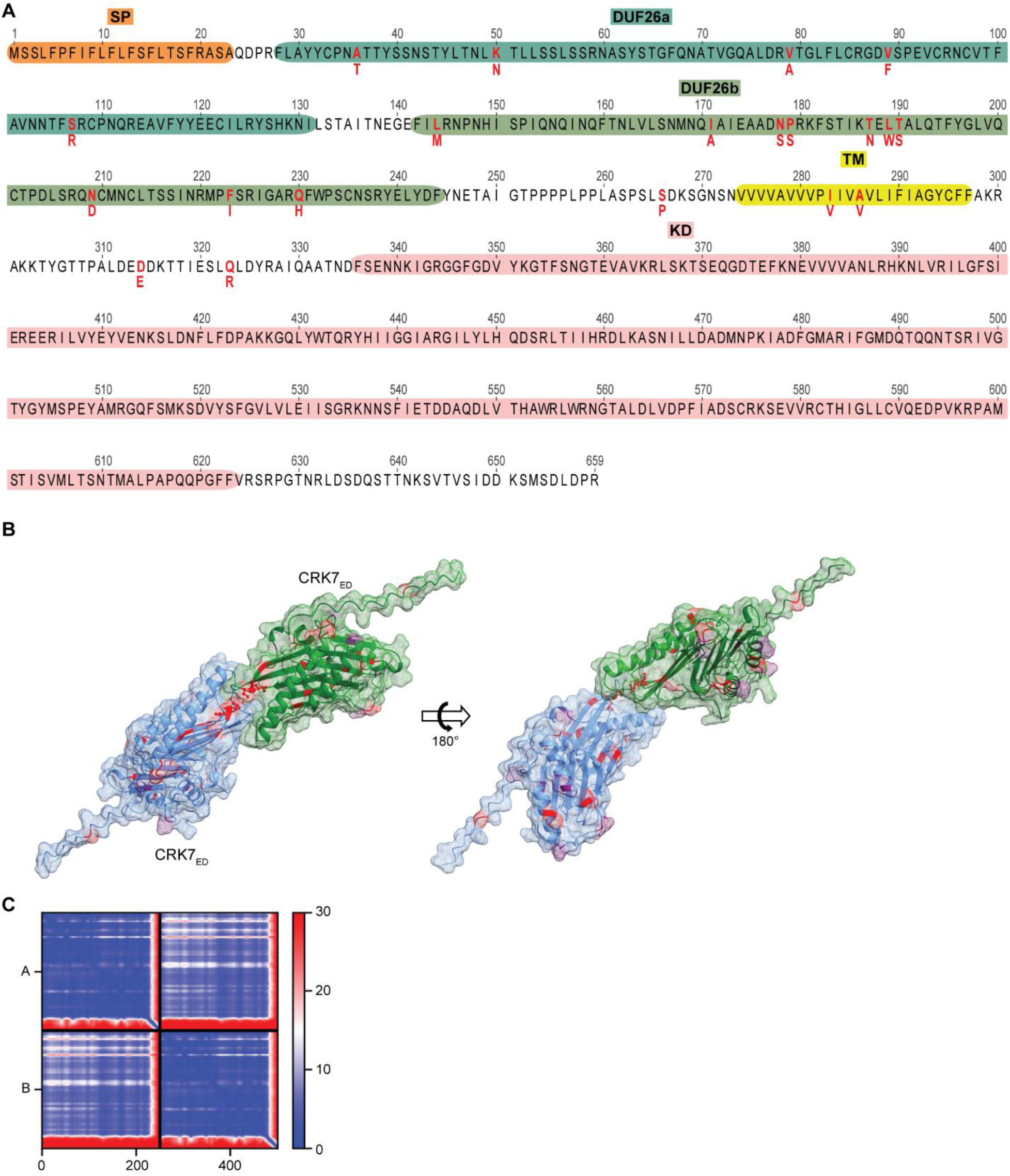
Conserved amino acid exchanges in WTA-insensitive *Arabidopsis* accessions accumulate in the extracellular domain of CRK7. (**A**) Amino acid sequence of the Col-0 CRK7 protein. Red letters indicate conserved amino acid exchanges in WTA-insensitive *Arabidopsis thaliana* accessions. SP: signal peptide, TM: transmembrane domain, KD: kinase domain (**B**) AlphaFold (*30, 31*) prediction of CRK7-ectodomain homodimer. Predicted N-glycosylation sites are highlighted in purple. Red amino acids represent amino acids altered in WTA-insensitive *Arabidopsis thaliana* accessions. (**C**) Predicted Alignment Error (PAE) of the CRK7-ectodomain homodimer. Blue color represents high prediction confidence, red color represents low prediction confidence.

**Fig. S14.**
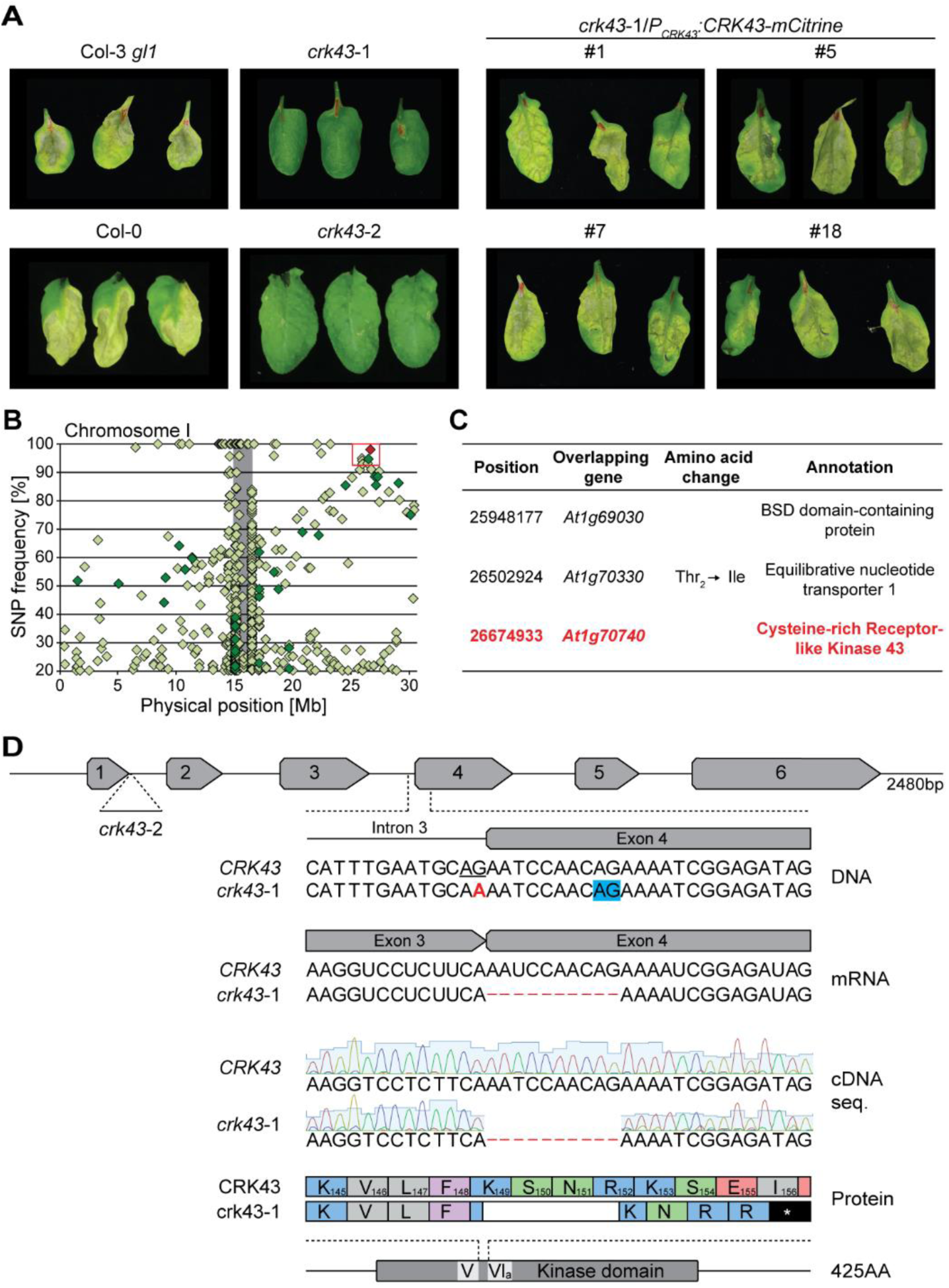
CRK43 is required for WTA-induced cell death. (**A**) WTA infiltration of indicated mutants and corresponding wild types (Col-0 for T-DNA lines, Col-3 *gl1* for *crk7*-4), together with 4 independent complementation lines expressing mCitrine-tagged CRK43 under control of the native promoter in the *crk43*-2 background. Leaves were infiltrated with 1 µg/ml WTA in 10 mM MgCl_2_. Pictures show three infiltrated leaves of three individual plants at 6 dpi. (**B**) Identification of the *crk43*-2 mutant using mapping by sequencing of F2 plants with a suppressor phenotype. Diamonds indicate the position and frequency of the synonymous (light-green) and nonsynonymous (dark-green) single-nucleotide polymorphisms (SNPs). Candidate SNPs are highlighted by the red box. The red diamond indicates the *crk43*-2 mutation. Grey background indicates centromere regions. (**C**) Candidate suppressor SNPs identified on Chromosome I via mapping-by-sequencing. (**D**) Schematic representation of the *CRK43* gene on Chromosome I of *Arabidopsis* Col-0. The T-DNA insertion loci of *crk43*-2 is depicted, as well as the G to A mutation (red font) in the *crk43*-1 mutant. The *crk43*-1 mutation converts the canonical “AG” splice acceptor (*69*) at the end of intron 3 (underlined) into “AA.” cDNA analysis from Col-0 and *crk43*-1 revealed that all analysed suppressor lines carry a 10 bp deletion in exon 4, consistent with the spliceosome utilizing the next downstream “AG” as an alternative acceptor (blue box). The deletion introduces a frameshift and premature stop codon between kinase subdomains V and Via (*70*), resulting in a truncated protein lacking over half of the kinase domain.

**Fig. S15.**
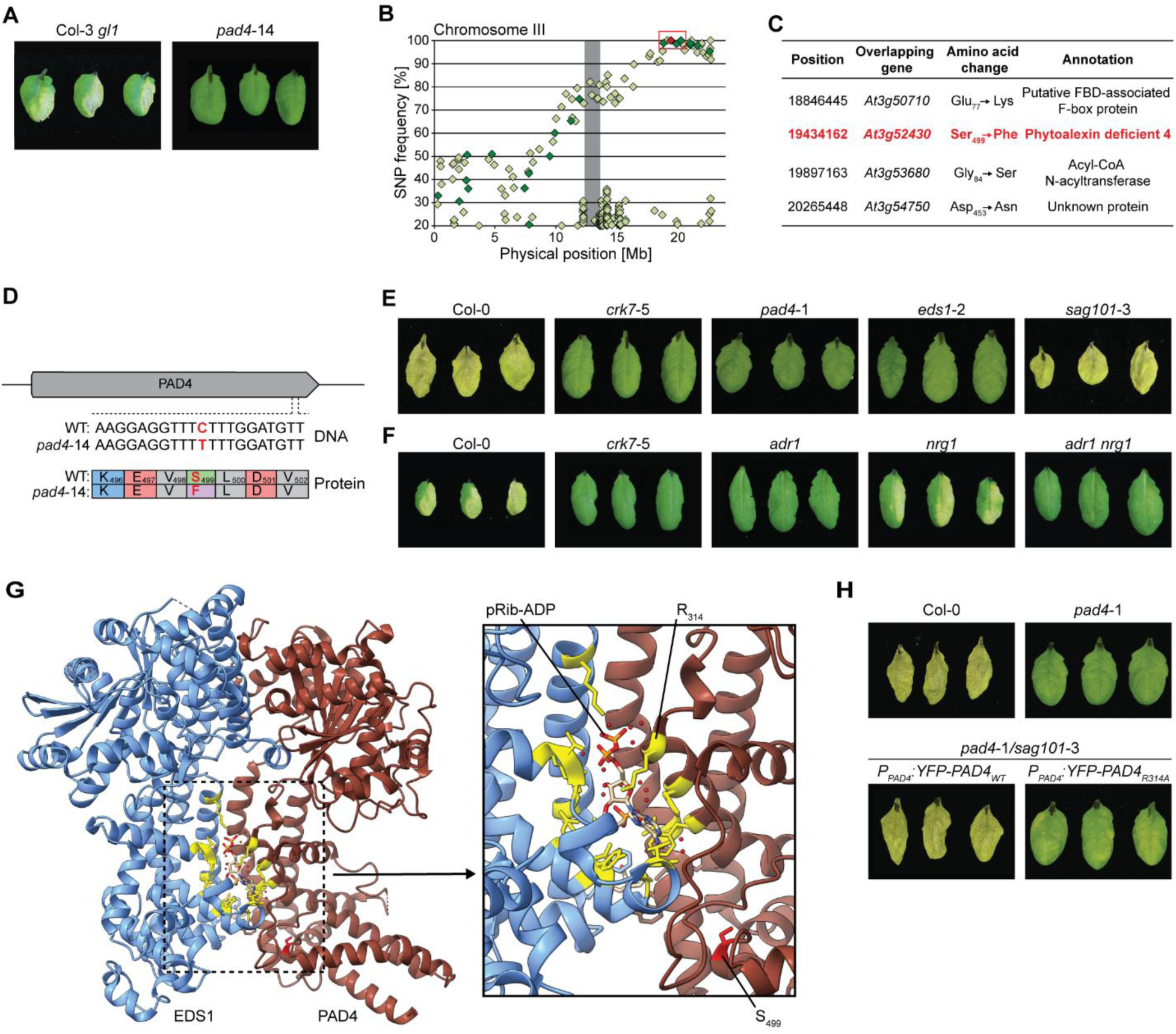
WTA-induced cell death requires the EDS1-PAD4-ADR1 signaling hub. (**A**) WTA infiltration of the *pad4*-14 mutant and the corresponding wild-type Col-3 *gl1*. Leaves were infiltrated with 1 µg/ml WTA in 10 mM MgCl_2_. Pictures show three infiltrated leaves of three individual plants at 6 dpi. (**B**) Identification of the *pad4*-14 mutant using mapping by sequencing of F2 plants with a suppressor phenotype. Diamonds indicate the position and frequency of the synonymous (light-green) and nonsynonymous (dark-green) single-nucleotide polymorphisms (SNPs). Candidate SNPs are highlighted by the red box. The red diamond indicates the *pad4*-14 mutation. Grey background indicates the centromere region. (**C**) Candidate suppressor SNPs identified on Chromosome III via mapping-by-sequencing. (**D**) Schematic representation of the *PAD4* gene on Chromosome III of *Arabidopsis thaliana* Col-0. The amino acid exchange resulting from the *pad4*-14 mutation is depicted. (**E** and **F**) Infiltration of *eds1-2*, *pad4-1*, and *sag101*-3 (E) as well as *adr1* triple, *nrg1* triple, and *adr1 nrg1* hextuple (F) mutants. Col-0 and *crk7*-5 were used as positive and negative control, respectively. Leaves were infiltrated with 1 µg/ml WTA in 10 mM MgCl_2_. Pictures show three infiltrated leaves of three individual plants at 6 dpi. (**G**) Protein structure of the EDS1-(blue) PAD4 (dark-red) heterodimer bound to pRib-ADP (*71*). The EDS1 and PAD4 amino acids involved in pRib-ADP binding are depicted in yellow. Amino acid Serine_499_, which is exchanged to a Phenylalanine in the *pad4*-14 mutant, is depicted in red. (**H**) Infiltration of *pad4*-1/*sag101*-3 mutant plants complemented with wild-type PAD4 and a mutant allele of PAD4 with an amino acid exchange (R_314_A) in the pRib-ADP binding site, with 1 µg/ml WTA. Col-0 and the *pad4*-1 mutant were used as controls.

**Table S1:**
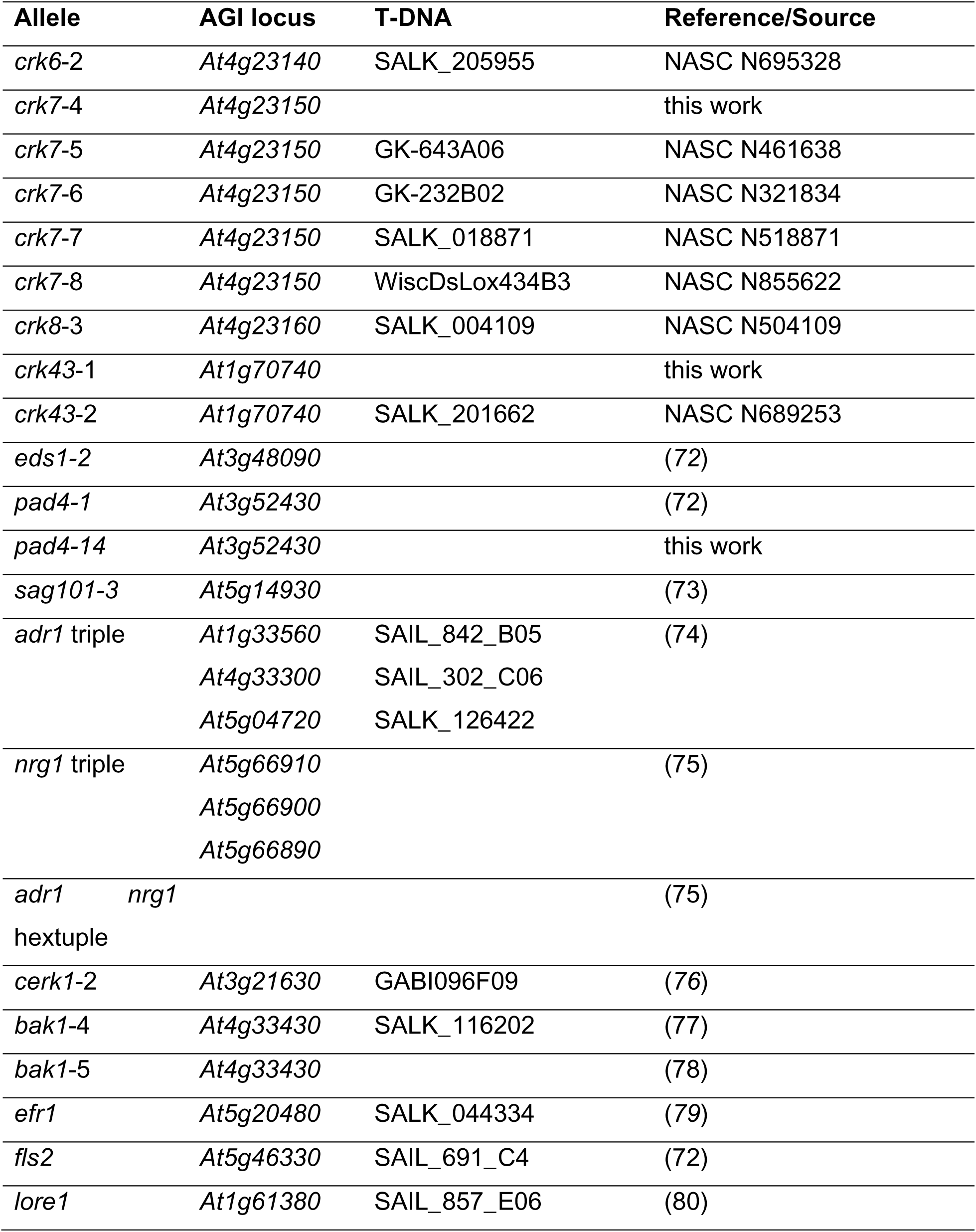
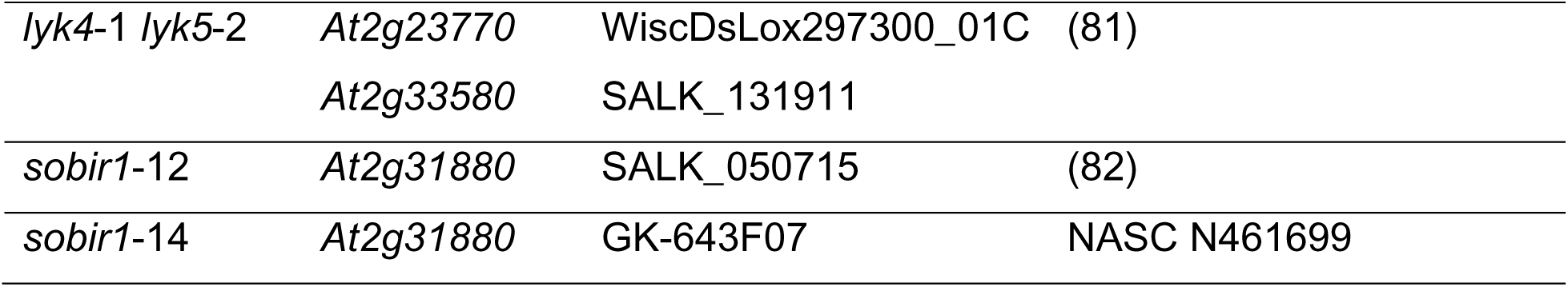
List of *Arabidopsis* mutants used in this work.

**Table S2:**
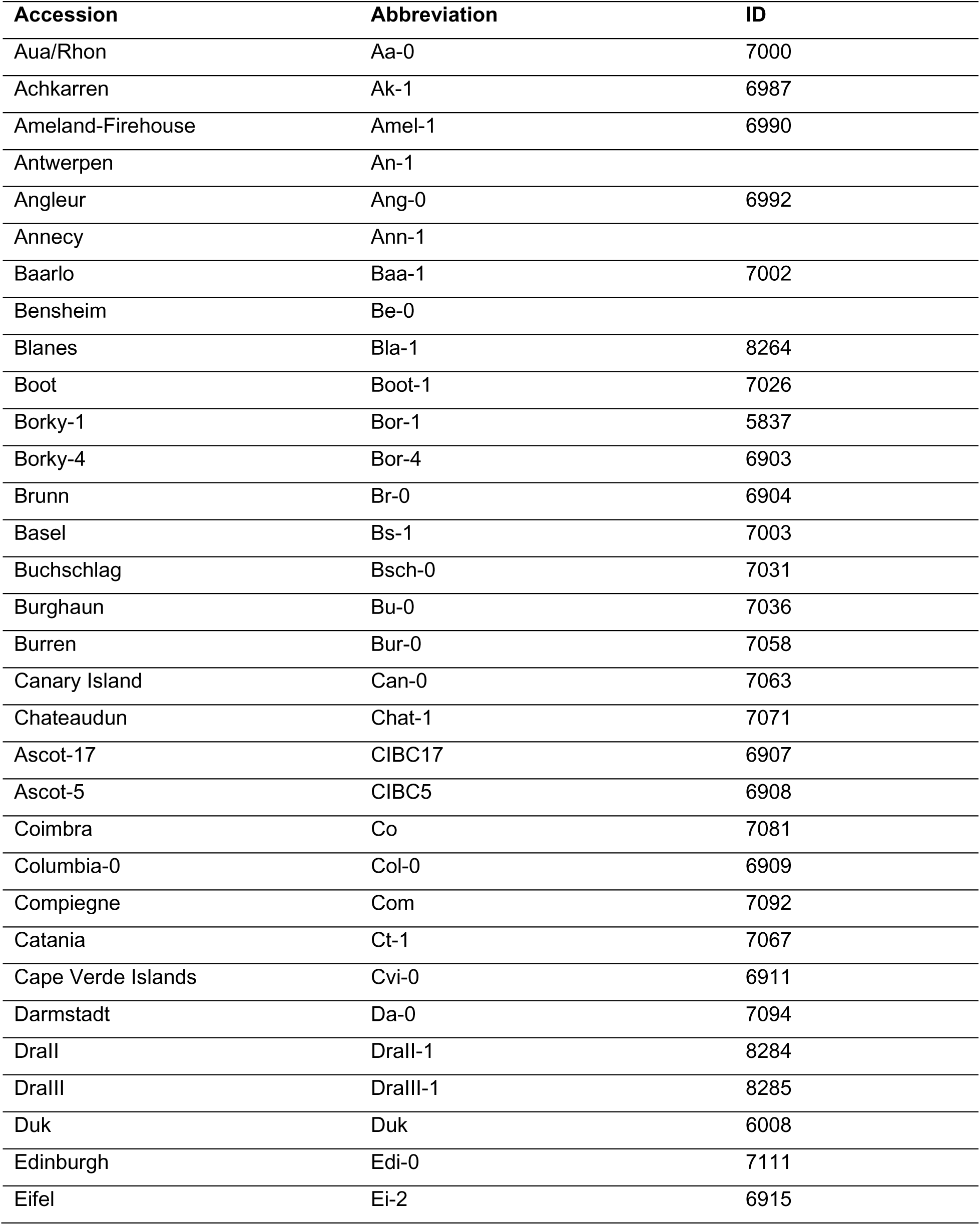
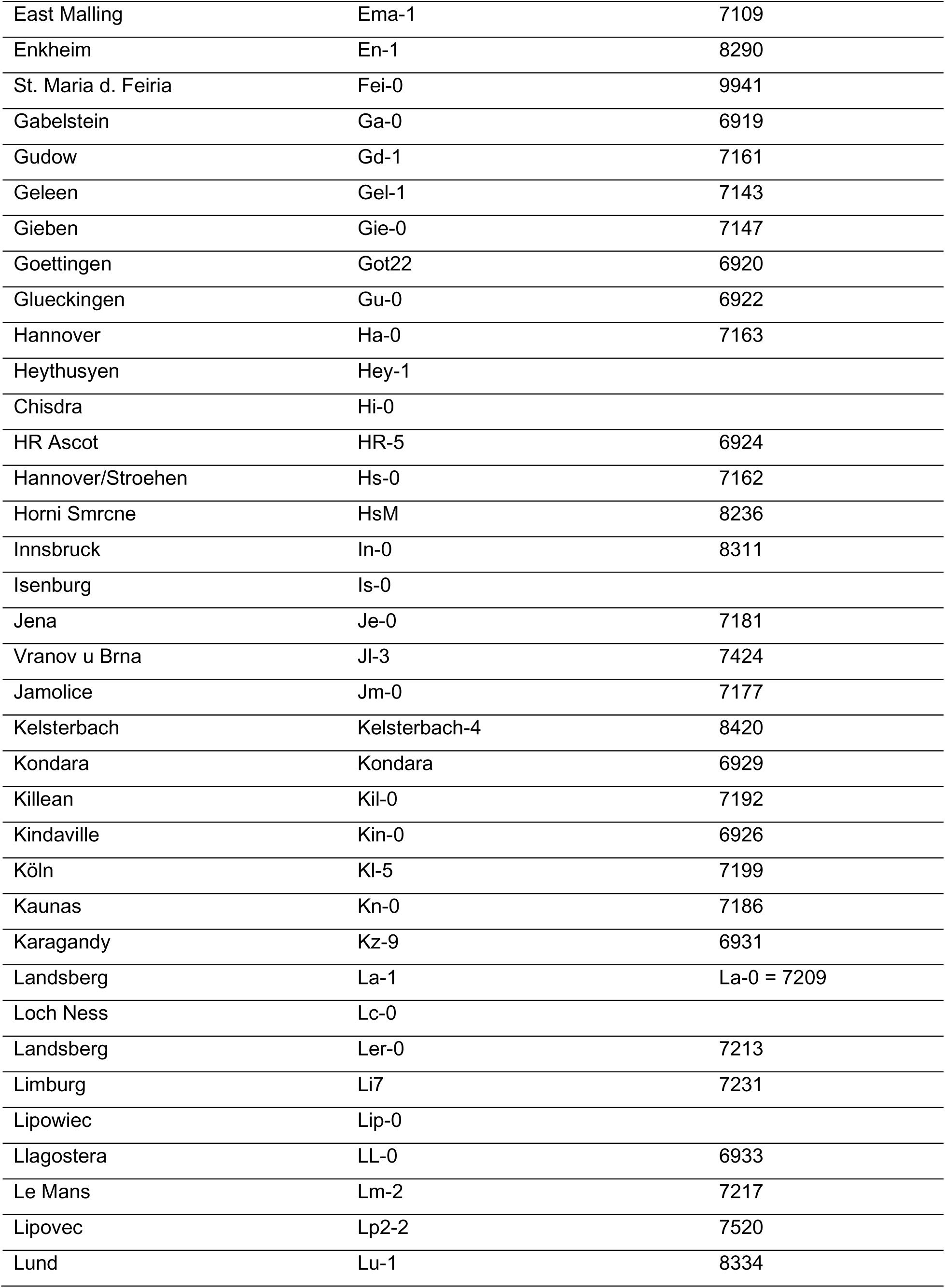
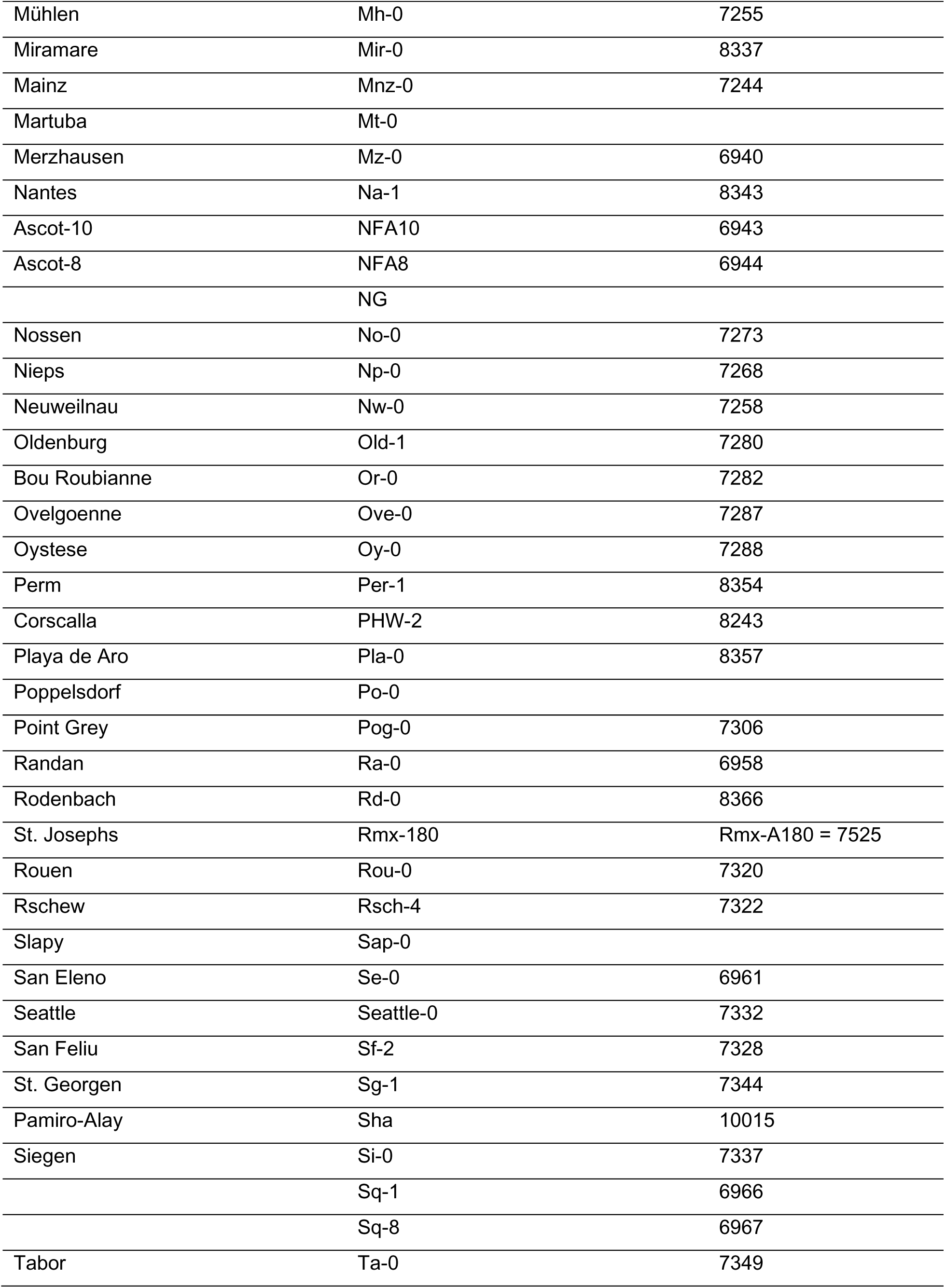
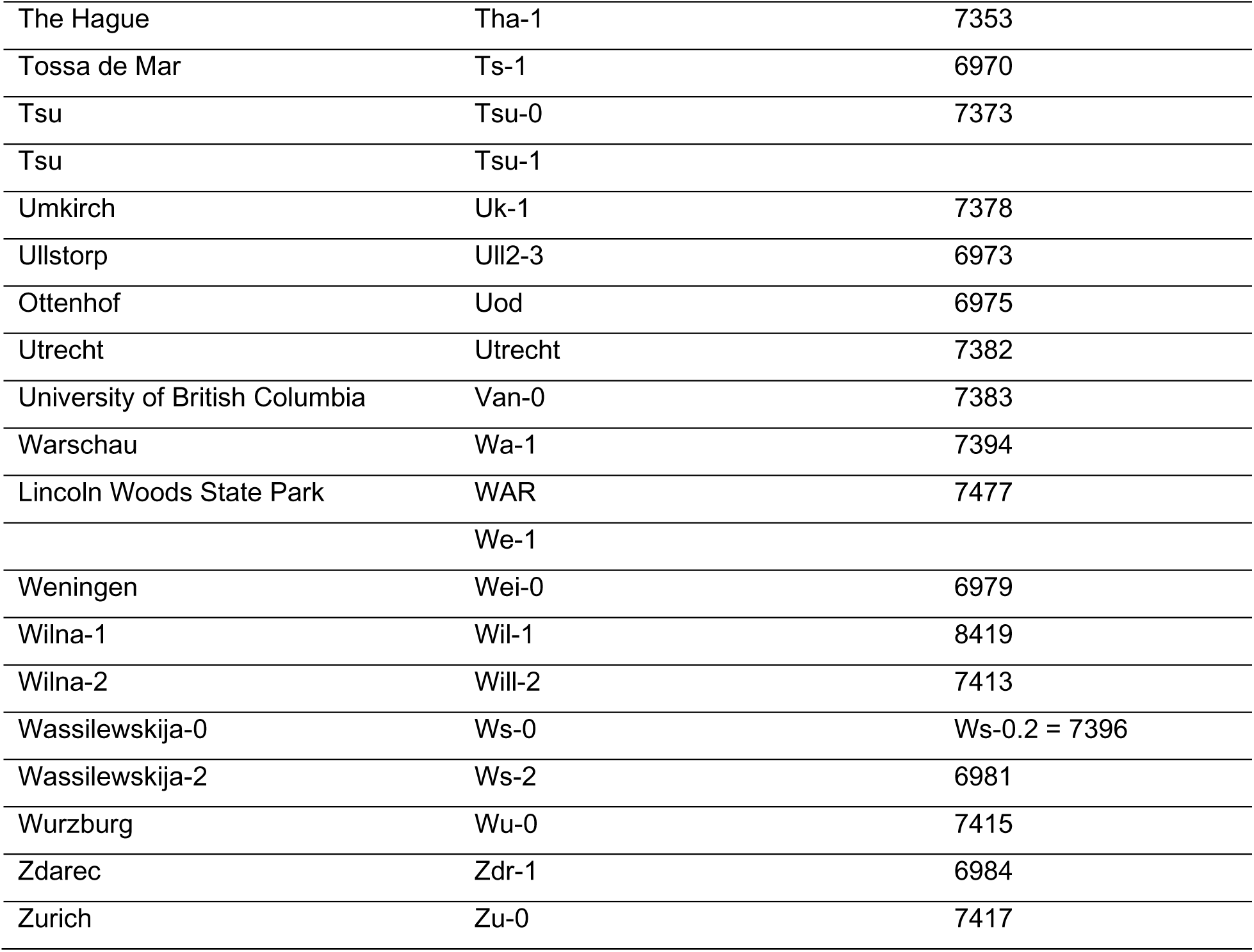
*Arabidopsis* Ecotypes used in this work. ID: 1001 Genomes.org (*29*) Accession ID

**Table S3:**
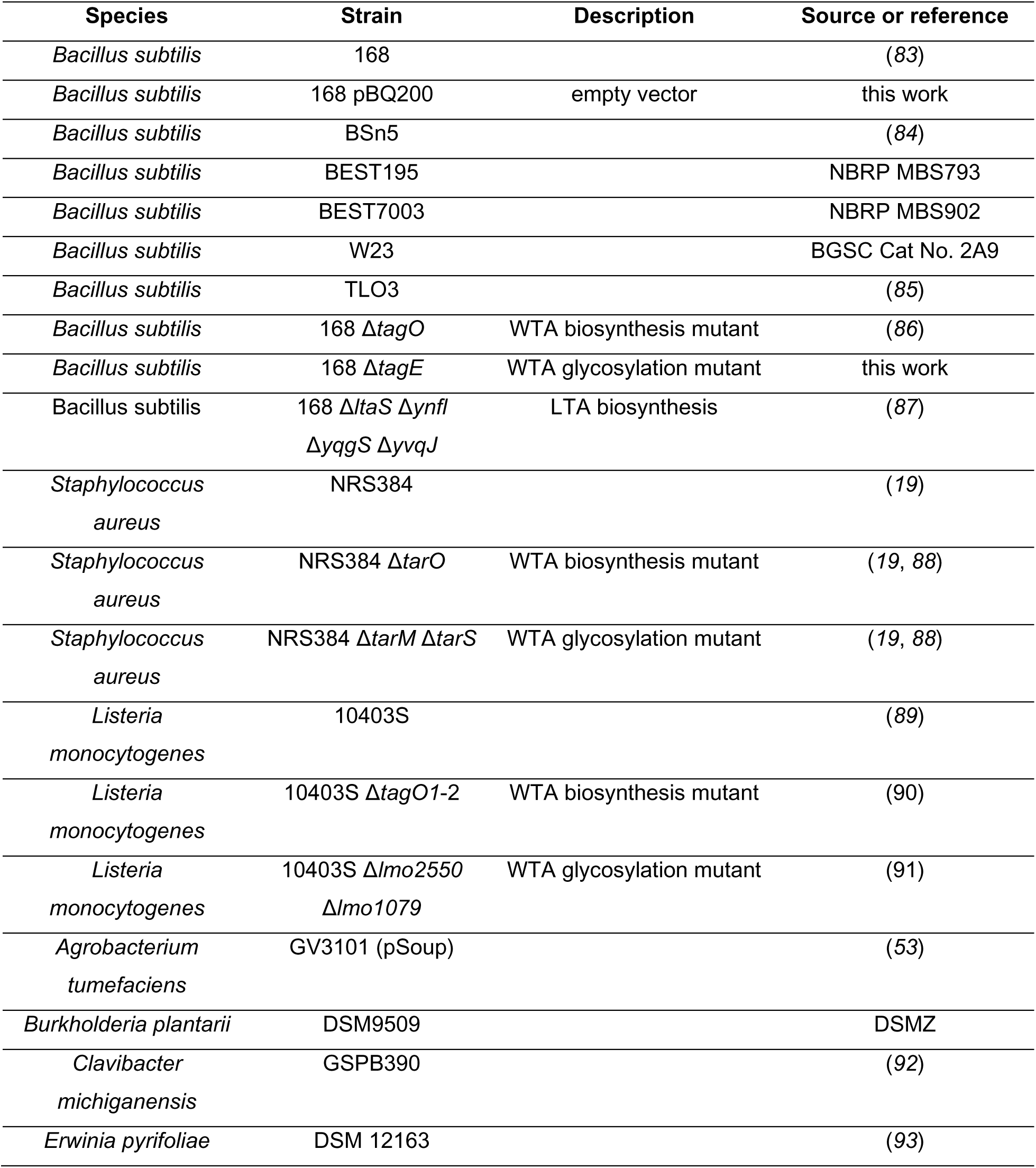
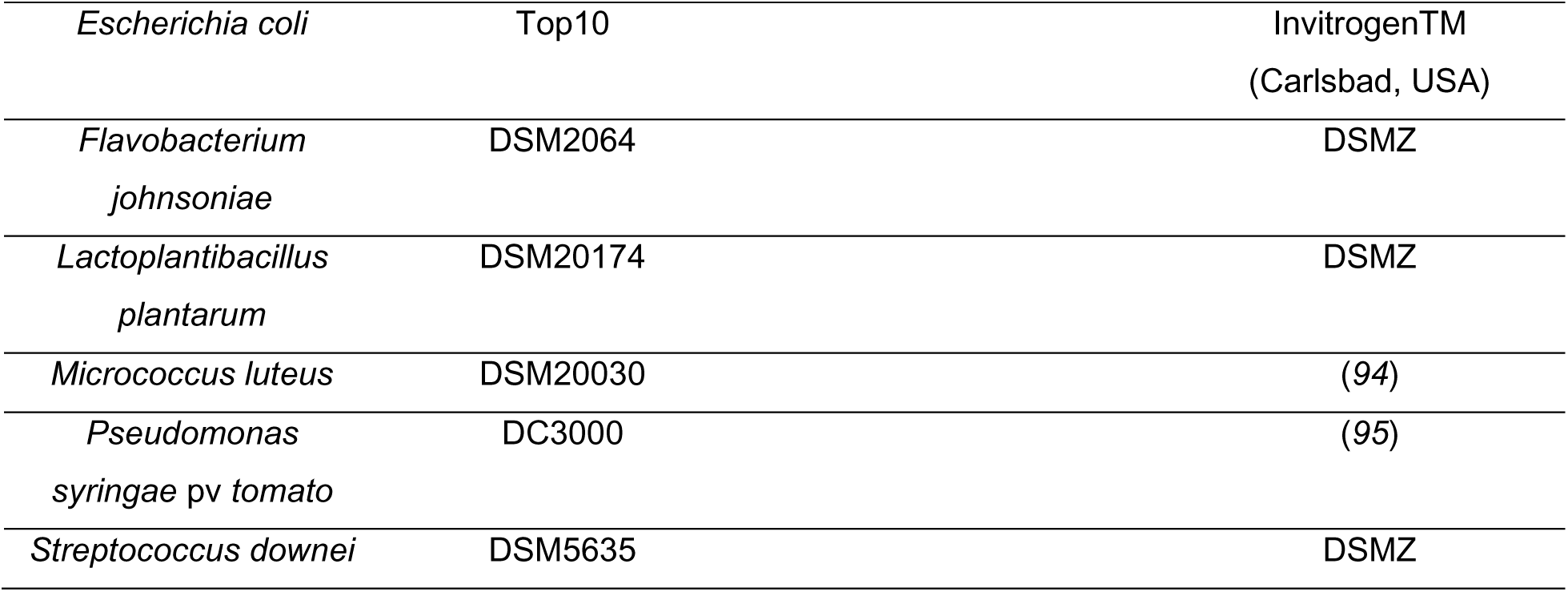
Bacterial species and strains used in this work. NBRP: National BioResource Project (Shizuoka, Japan); BGSC: Bacillus Genetic Stock Center (Columbus, USA); DSMZ: German Collection of Microorganisms and Cell Cultures (Braunschweig, Germany)

**Table S4:**
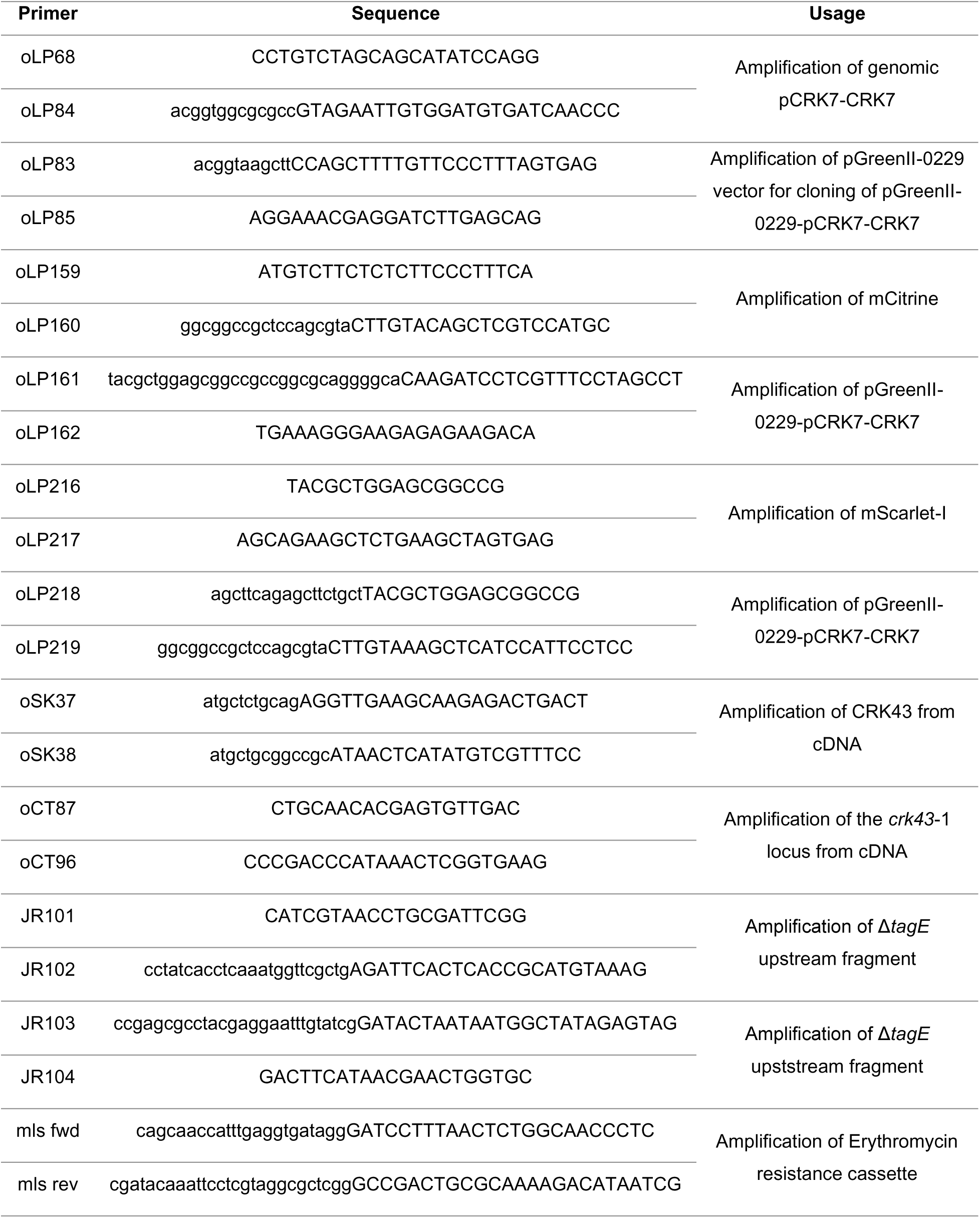
Primers used in this work. Capital letters: Primer binding site. Lowercase letters: 5’ extension

**Table S5:**
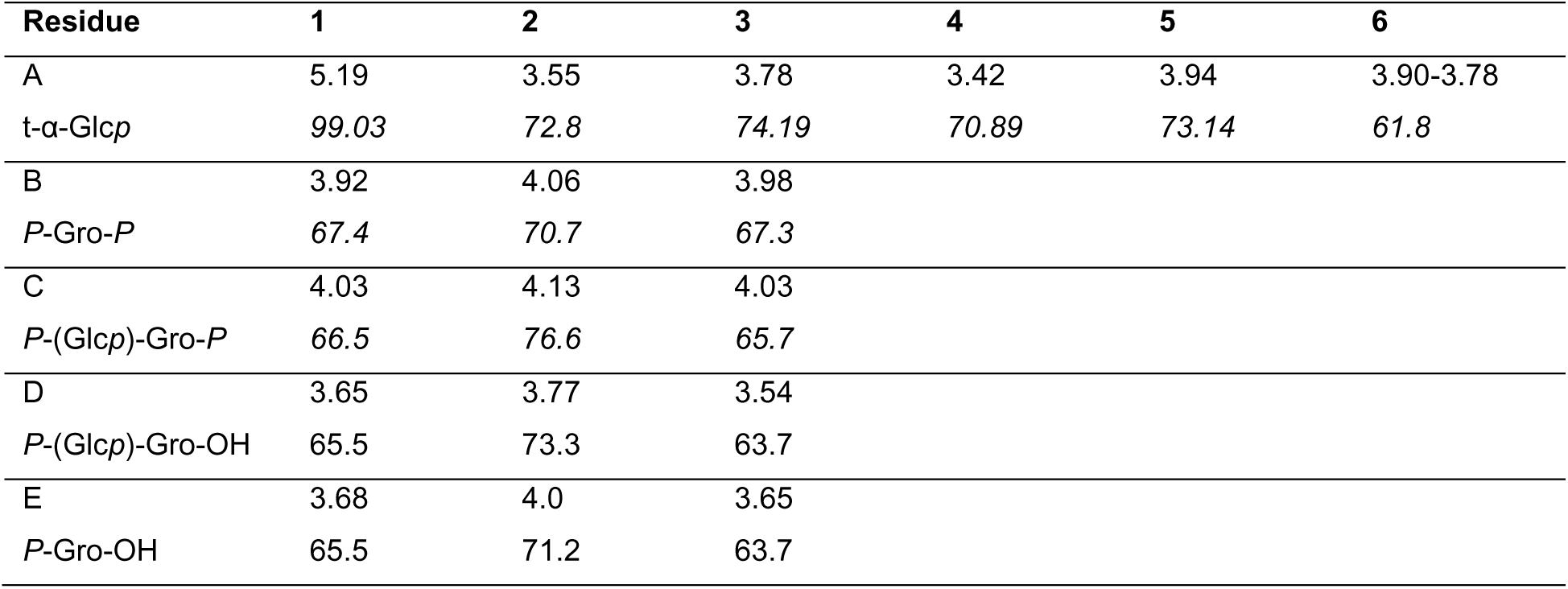
NMR chemical shifts of the *B*. *subtilis* glycan-1, measured at 600 MHz, 298 K, in D_2_O.

## Notes

### Competing Interest Statement

The authors have declared no competing interest.

